# Collective cell sorting requires contractile cortical waves in germline cells

**DOI:** 10.1101/2020.01.06.895847

**Authors:** Soline Chanet, Jean-René Huynh

## Abstract

Encapsulation of germline cells by layers of somatic cells forms the basic unit of female reproduction called primordial follicles in mammals and egg chambers in *Drosophila*. How germline and somatic tissues are coordinated for the morphogenesis of each separated unit remains poorly understood. Here, using improved live-imaging of *Drosophila* ovaries, we uncovered periodic actomyosin waves at the cortex of germ cells. These contractile waves are associated with pressure release blebs, which project from germ cells into somatic cells. We demonstrate that these cortical activities, together with cadherin-based adhesion, are required to sort each germline cyst as one collective unit. Genetic perturbations of cortical contractility, blebs protrusion or adhesion between germline and somatic cells induced failures to encapsulate any germ cells or the inclusion of too many germ cells or even the mechanical split of germline cysts. Our results reveal that germ cells play an active role in the physical coupling with somatic cells to produce the female gamete.

## INTRODUCTION

During development, tissues from different origins often cooperate and coordinate their morphogenetic movements to generate complex organs. Formation of the female gamete for example requires tight coordination between germ cells and the surrounding somatic tissue. During their differentiation, germ cells undergo several rounds of mitosis before entering meiosis. In most species, these mitoses are incomplete giving rise to cysts of cells interconnected by cytoplasmic bridges (Pepling et al., 1999). Each germline cyst is then surrounded by cells of somatic origin called pre-granulosa cells in mammals and follicle cells in *Drosophila* (Elkouby and Mullins, 2017). In mammals, pre-granulosa cells invade in between germ cells and each cyst eventually breaks down (CBD) into single cells encased by granulosa cells, forming primordial follicles (PFs) (Lei and Spradling, 2013). In *Drosophila*, egg chambers are made of precisely 16 germ cells surrounded by an epithelium of follicle cells (De Cuevas et al., 1997). Follicle cells do not invade the germline and intercellular bridges are maintained throughout oogenesis. Encapsulation is a conserved and important process to investigate as PFs and egg chambers give rise to the future female gamete, and defects in this step lead to sterility. Studies on the role of somatic cells during this step revealed general principles of epithelium morphogenesis (Godt and Tepass, 2003; Horne-Badovinac and Bilder, 2005; Sarpal et al., 2012; St Johnston and Sanson, 2011). In contrast, the contribution of germ cells to encapsulation remains poorly characterized, and germ cells are assumed to be passive and transported by somatic cells.

In *Drosophila*, these early steps of oogenesis take place in a specialized structure called the germarium at the anterior tip of the ovary (Figure 1a) (Huynh and St Johnston, 2004). The germarium contains both germline and somatic stem cells, which divide to produce egg chambers throughout adult life. Germline stem cells (GSCs) are located at the most anterior tip of the region 1 of the germarium and give rise to cystoblasts. Cystoblasts then undergo four rounds of incomplete mitosis to generate cysts of 16 cells interconnected by ring canals. Only one cell per cyst becomes an oocyte and completes meiosis. The remaining 15 cells differentiate as nurse cells and synthesize nutrients and RNAs required for oocyte growth and maturation. Once made of 16 cells, germline cysts enter region 2a, where they come into contact with follicle cells (FCs) of somatic origin produced by a population of follicle stem cells (FSCs) (Reilein et al., 2017). In region 2a, germline cysts are round, and several cysts can be found at similar stages of development. Then, only one cyst at a time moves to region 2b and flattens to take the shape of a disc spanning the width of the germarium. All 16 cells of each cyst move collectively as a unit. It is during this transition from region 2a to 2b, that encapsulation starts. The 16 cells of each cyst become separated from other cysts by ingressing FCs that migrate centripetally (Horne-Badovinac and Bilder, 2005; Morris and Spradling, 2011). Moving posteriorly to region 3 (aka stage 1), cysts increase in volume and become round again, encased by a monolayer of around 30 epithelial FCs. FCs at both poles of the cyst intercalate and form stalks of cells, which pinch off the newly formed egg chamber from the germarium into the vitellarium (Morris and Spradling, 2011). In the vitellarium, egg chambers grow rapidly as separate units and polarize to become competent for fertilization.

**Figure 1:**
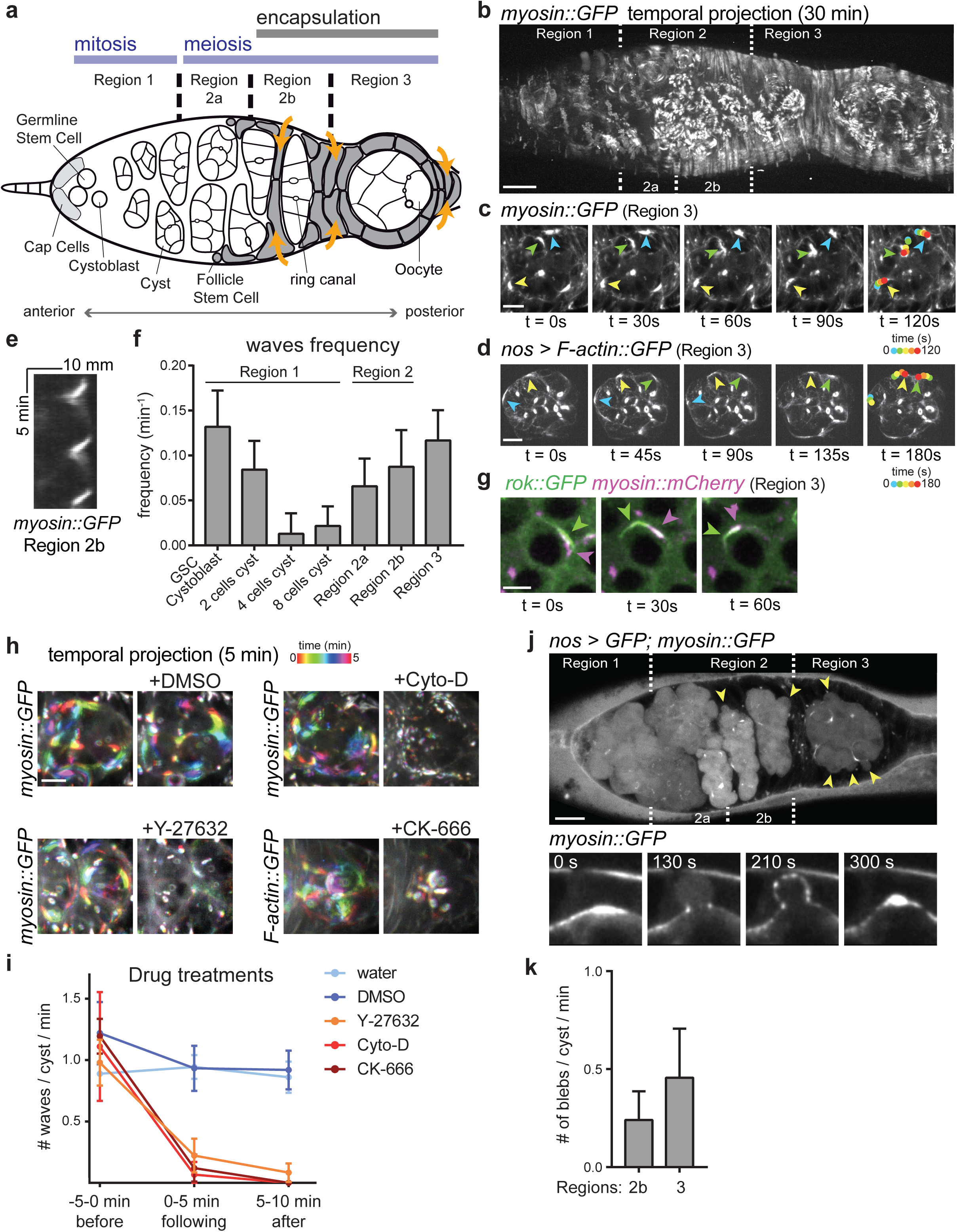
Germ cells generate actomyosin traveling waves associated with pressure released blebs. (**a**) Scheme of a germarium. The germarium is divided into four morphological regions along the anterior–posterior axis. At the anterior tip, the germline stem cell and its progeny divide to produce 16-cell germline cysts that are connected by ring canals. The cysts then enter meiosis. At the transition between region 2a and 2b resides follicle stem cells. They produce follicle cells (gray) that surround and separate germline cysts (orange arrows) in a process called encapsulation. In region 3 the cyst adopts the shape of a sphere and is surrounded by one sheet of somatic cells, the newly formed egg chamber will be progressively pinch out of the germarium. Anterior is on the left, posterior on the right. (**b**) Temporal projection of a 30 min movie of a germarium expressing *sqh::GFP* that stains myosin. (**c**) Time-laps images of *sqh::GFP* signal showing myosin travelling waves around the cell cortexes of a germline cyst in region 3. On the last image, waves displacement is represented with time-colored dots. (**d**) Time-laps images of *LifeAct::GFP* signal that stains F-actin expressed specifically in the germline showing actin travelling waves around the cell cortexes of a germline cyst in region 3. On the last image, waves displacement is represented with time-colored dots. (**e**) Kymograph showing three consecutive waves of *sqh::GFP* (myosin) around the cortex of one cell from a cyst in region 2b. (**f**) Quantification of waves frequency depending on the developmental time of germline cyst cells. n = 6 GSC or Cystoblast, n = 12 cells of 2-cell cysts, n = 21 cells of 4-cell cysts, n = 33 cells of 8-cell cysts, n = 61 cells of region 2a cysts, n = 45 cells of region 2b cysts, n = 42 cells of region 3 cysts; 4 germaria. (**g**) Time-laps images of a germ cell expressing both *rok::GFP* (green) and *sqh::mCherry* (myosin, magenta). Green and magenta arrows point to the position (middle) of rok and myosin waves respectively. (**h**) Temporal projection of 5min movies of germline cysts in region 3 before (left) or after (right) drug or vehicle addition to the medium as indicated. Time is color-coded, such that rainbow signal indicates a travelling wave. Myosin (*sqh::GFP*) or F-actin (*utr::GFP*) waves are examined. (**i**) Quantification of waves frequency per cyst in region 2b or 3 before and after drug or vehicle treatments. Mean and standard deviations (SD) are shown. n = 4 germaria treated with water, n = 5 germaria treated with DMSO, n = 6 germaria treated with Y27632, n = 6 germaria treated with Cyto-D, n = 5 germaria treated with CK-666. (**j**, top) cytoplasmic expression of GFP in the germline reveals the presence of bleb protrusions (arrows). (bottom) still-images following a bleb formation and retraction, *sqh::GFP* stains myosin. (**k**) Quantification of bleb frequency per cysts in region 2b and region 3. Mean and SD are shown. n = 19 region 2b cysts, n = 13 region 3 cysts, 19 germaria. Scale bars, 10μm (b, j), 5μm (c, d, g, h). See also Movies 1-6 and Supplementary Figure 1.

How the morphogenetic movements of encapsulation are coordinated between germ cells and somatic cells is not known. Mechanical forces that shape cells and tissue are usually produced by actomyosin contractility where the molecular motor myosin 2 contract cortical filamentous actin (F-actin). To be effective, these contractile forces must be transmitted to the extracellular substrate or neighboring cells through links between F-actin network and junctional complexes, such as cadherins and integrins (Lecuit et al., 2011; Salbreux et al., 2012). In both mammals and flies, cadherins rather than integrins mediate interactions between somatic and germ cells at these early stages (Bendel-Stenzel et al., 2000; Godt and Tepass, 2003). In particular, in flies, E-Cadherin forms a gradient in both FCs and germ cells to help position the oocyte at the posterior of the egg chamber (Becam et al., 2005; Godt and Tepass, 1998; Gonzalez-Reyes and St Johnston, 1998). β-catenin (*armadillo*, *arm* in *Drosophila*) and α-catenin link E-Cadherin to the underlying actomyosin cortex (Peifer et al., 1993; Sarpal et al., 2012; White et al., 1998).

In this study, we used hydrogel-based live imaging, genetics and image analyses to investigate the role of germ cells during the very first steps of encapsulation. We show that germline cysts actively generate forces both to maintain their position in the germarium while being surrounded by FCs, and to preserve their integrity as groups of 16 cells.

## RESULTS

### 1) Germ cells generate actomyosin traveling waves

To investigate a potential role of germ cells during encapsulation, we first looked at actomyosin dynamics in the germline. To monitor the dynamics of the actomyosin cytoskeleton, we used a GFP-tagged version of the regulatory light chain of the non-muscle myosin 2 (sqh::GFP, and hereafter referred as myosin::GFP) expressed under its own promoter (Royou et al., 2002), and the actin reporters, LifeAct (that we expressed exclusively in the germline) or Utrophin (Utr), expressed ubiquitously (Huelsmann et al., 2013; Rauzi et al., 2010). Strikingly, we observed waves of myosin and F-actin at the cortex of germline cysts (Figure 1b-e, Movies 1-2). The frequency of these waves was not uniform across the germarium (Figure 1b, f). Wave frequency was high in GSCs with a low myosin intensity signal, and then faded away in the mitotic region. It became high again, and both more periodic and intense in region 2a/2b. Interestingly, this increase in region 2a/2b corresponds to the first contacts between somatic follicle cells and germline cysts (Figure 1b, f). These results indicated that the cortical dynamics of the germ cells vary according to cyst differentiation. In region 2b and region 3, we measured a period of the waves of 13,5 +/-1.2 min and 9.4 +/-0.5 min respectively. We found that *sqh::Dendra2* photoconverted on one side of a cell could travel to the opposite side demonstrating that myosin was travelling (Figure S1a) (Roubinet et al., 2017). We measured an average speed of 0.028 +/− 0.0022 µm.s^-1^.

We next asked what regulates actomyosin waves. Cortical dynamics can be regulated by myosin activity, localization, F-actin network conformation (branched *vs.* linear) as well as polymerization and turn-over rates. Myosin activation often occurs downstream of RhoA and its effector, the myosin-activating kinase, Rho-associated coiled-coil kinase ROCK (*rok* in *Drosophila*) (Jaffe and Hall, 2005; Winter et al., 2001). We observed that ROCK also travelled as waves at the cortex of germ cells, but slightly ahead of myosin waves (Figure 1g, the time shift between ROCK and myosin waves was about 30s). To test if the activity of ROCK was required for the propagation of these waves, we made use of the chemical inhibitor Y27632. We used a water-based environment to be able to add this drug while recording. We adapted a live-imaging set up based on a photo-crosslinked PEG hydrogel, which could gently immobilize and maintain germarium morphology, while allowing diffusion of aqueous medium and drug treatments (Figure S1b) (Burnett et al., 2018). The addition of Y27632 stopped waves propagation within 1 to 2 min, indicating that ROCK activity was required for waves propagation (Figure 1h, i, Movie 3). As a control experiment, addition of water alone had no effect (Figure 1i).

Next, we tested the requirement for F-actin dynamics. We found that adding Cytochalasin D (Cyto-D), an inhibitor of actin turn-over, or CK-666, which inhibits Arp2/3 and actin branching, completely repressed waves propagation. In contrast, adding DMSO alone had no effect (Figure 1h, i, Movie 3). Finally, we found that addition of Colcemid, an inhibitor of microtubules polymerization, had no detectable effect on actomyosin waves dynamics (Movie 4).

Collectively, these results describe periodic oscillations of the actomyosin cytoskeleton at the cortex of germline cells; and demonstrate that waves propagation requires myosin activity and actin polymerization.

### 2) High cortical contractility in germ cells is associated with pressure release blebs

Strong contractions of the actomyosin network can induce ruptures of the cortex or its detachment from the plasma membrane (Charras and Paluch, 2008; Diz-Muñoz et al., 2013). These ruptures lead to the formation of cytoplasmic protrusions, called blebs, which release cytoplasmic hydrostatic pressure. Blebs are thus signs of strong cortical contractility. We often observed that following a wave of myosin, a break in the actomyosin meshwork formed and allowed the expansion of bleb protrusions deep into the somatic cell layers. Blebs expansion left only a ring of myosin at the neck (Figure 1j, Movie 5). The actomyosin meshwork then reformed inside the protrusion driving bleb retraction. Inhibiting cortical contractility by adding Cyto-D immediately eliminated blebs (Movie 6). This showed that blebs formation in germ cells was dependent on actomyosin contractility. We found that blebs frequency followed the increase in wave occurrences from region 2b to region 3 (Figure 1k). Noticeably, no blebbing was detected in region 1 of the germarium when cortical contractility is weak.

We concluded that oscillations of actomyosin at the germ cells cortex were contractile and associated with blebs. This high contractility in region 2b/3 suggested that germline cells could play an active role in the encapsulation process.

### 3) Alterations of cortical contractility in germ cells lead to the packaging of abnormal numbers of germ cells

In order to test the functional significance of contraction waves, we reduced cortical contractility in the germline, either by knocking down *chickadee* (*chic-RNAi*), the *Drosophila* homolog of profilin, required for actin polymerization; or *zipper* (*zip-RNAi*), which encodes for myosin 2 heavy chain. We achieved spatial and temporal specificity using either the *nanos*-Gal4 or *bam*-Gal4 driver, which are only expressed in germline cells. The *bam*-Gal4 driver is weaker than *nanos*-Gal4, but allows knocking down genes in region 2a to region 3 of the germarium without interfering with stem cells and the formation of germline cysts. Depleting *chic* or *zip* in germ cells effectively reduced cortical myosin intensity and waves frequency compared to a control knocked-down (*ctl-RNAi*) (Figure 2a, b, Movie 7). We obtained similar reduction of cortical contractility in germline cysts mutant for ROCK (*rok^2^*) (Figure S2a). We also tested the consequences of increasing cortical contractility by inhibiting *mbs*, the myosin binding subunit of the myosin phosphatase, which dephosphorylates and inhibits myosin. Depleting *mbs* in the germline (*mbs-RNAi*), increased waves frequency (Figure 2a, b, Movie 7).

**Figure 2:**
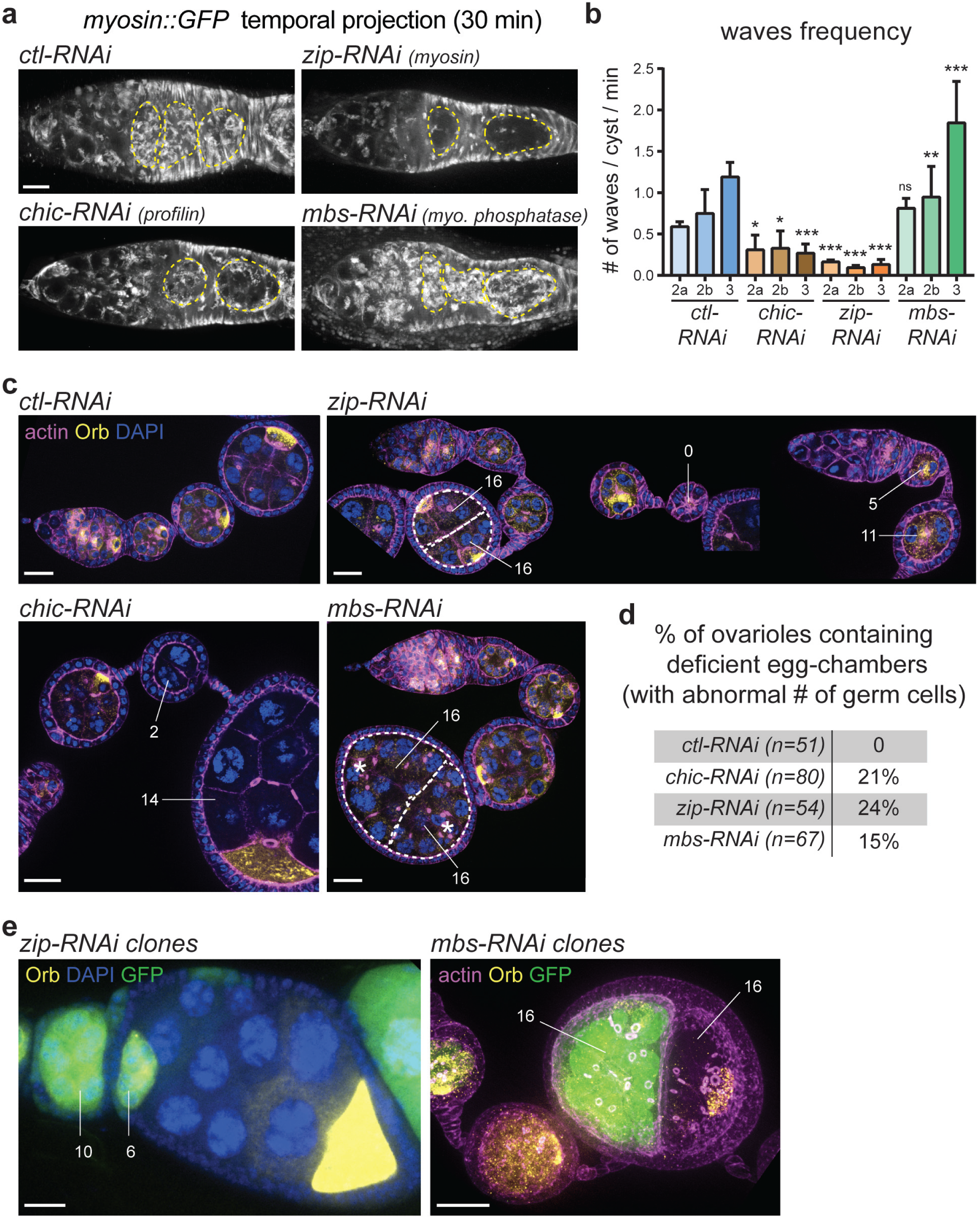
Alteration of germline contractility induces the formation of abnormal egg chambers. (**a**) Temporal projection of 30 min movies of germaria expressing *sqh::GFP* to follow myosin waves in the indicated mutant conditions. Germlines cysts are outlined with yellow dotted lines. (**b**) Quantification of wave frequency per cyst in region 2 and 3 of the germarium. Indicated RNAi were expressed specifically in the germline. Mean and SD are shown. n = 5 cysts in region 2a, n = 8 cysts in region 2b, n = 8 cysts in region 3, 5 *ctl-RNAi* germaria; n = 5 cysts in region 2a, n = 4 cysts in region 2b, n = 5 cysts in region 3, 5 *chic-RNAi* germaria; n = 4 cysts in region 2a, n = 6 cysts in region 2b, n = 5 cysts in region 3, 3 *zip-RNAi* germaria; n = 5 cysts in region 2a, n = 9 cysts in region 2b, n = 7 cysts in region 3, 5 *mbs-RNAi* germaria. * *P* < 0.05, ** *P* < 0.01, \*\*\**P* < 0.001, ns, not significant, *t*-test. (**c**) Fixed images of ovarioles stained with phalloidin to mark actin (magenta), Orb that marks the oocyte (yellow) and DAPI that marks the DNA (blue). Depletion of *zip*, *chic* or *mbs* in the germline (*zip-RNAi, chic-RNAi, mbs-RNAi*) lead to the formation of egg chambers with abnormal number of germ cells. White dotted lines underline two different cysts packaged together in one egg chamber (both made of 16 cells). Numbers indicate the number of cells included in each unit. Asterisks mark the position of the two oocytes of the two *mbs-RNAi* cysts. (Note: *mbs-RNAi* induces smaller ring canals leading to dedifferentiation of the oocyte and progressive loss of Orb accumulation). (**d**) Quantification of the occurrence phenotypes (egg chambers made of abnormal of germ cells) in fixed ovarioles for the different mutant conditions. (**e**, left) fixed mosaic ovariole showing a GFP+ *zip-RNAi* cyst that is split between two egg chambers (numbers indicate the number of cells from the original cyst that have been distributed between the two egg chambers). (right) fixed mosaic ovariole showing one GFP+ *mbs-RNAi* cyst packaged with an unmarked wild-type cyst within the same egg chamber. Scale bars, 10μm (a), 20μm (c, e). See also Supplementary Figure 2 and Movies 7.

The most striking phenotype induced by both reducing or increasing contraction waves in the germline was an abnormal number of germ cells per egg chamber, detected during mid-oogenesis (12 to 36h after leaving the germarium) (Figure 2c, d). Instead of egg chambers containing 16 germ cells with one oocyte, mutant egg chambers were made of a number of germ cells ranging from 0 to 32 cells, with 0, 1 or 2 oocytes (Figure 2c, d, oocyte marked by Orb). The abnormal number of germ cells could come from defects in mitosis of single-cell precursors (abnormal number of divisions or defective abscission of GSCs/cystoblasts) (Hawkins et al., 1996; Mathieu et al., 2013); or defects in cell sorting, with the encapsulation of germ cells from different cysts into the same egg chamber. To distinguish between these two hypotheses, we performed a cell-lineage analysis using the FLP-out technique to label germline cysts generated by a single-cell precursor. In our experiment, RNAi expressing cysts were GFP+ and wild type cysts GFP-. We found that egg chambers containing 32 germ cells were made of two different 16-cell cysts packaged together, and not a single 32-cell cyst caused by abnormal divisions (Figure 2e). Similar results were obtained with germline clones mutant for *sqh^1^* (Figure S2b). We also found that egg chambers with less than 16 germ cells were associated with neighboring egg chambers containing the missing complement of GFP+ germ cells (Figure 2e, 10 GFP+ germ cells in one egg chamber are associated with the missing 6 GFP+ germ cells in the neighboring egg chamber). Long stalks of FCs correlated with pseudo-egg chamber empty of any germ cell (Figure 2c).

We concluded that altering cortical contractility induced defects in sorting germline cysts into groups of 16 cells and resulted in the formation of unfertile egg chambers with abnormal numbers of germ cells. It further suggested that the cause of these phenotypes could be encapsulation defects at earlier stages of oogenesis.

### 4) Abnormal number of germ cells in contractility mutants are caused by encapsulation defects

To test if abnormal egg chambers originated from encapsulation defects, we imaged live the early steps of encapsulation in hydrogel. We followed the displacement of individual cysts along the anterior-posterior (a-p) axis, at the time they started to be separated by FCs (in region 2b and 3 of the germarium). We used cap cells at the anterior tip of the germarium as a fixed reference point and measured cyst displacement by tracking their movements over 1 to 2h (Figure 3a, b, initial positions indicated by dotted lines and final positions with continuous line). In *ctl-RNAi* conditions, we observed that when somatic cells ingressed to separate two cysts, the older cyst was slightly displaced toward the posterior of the germarium (positive orientation), and the younger cyst was slightly displaced toward the anterior (negative orientation) (Figure 3b, Figure S3, Movie 8). Overall cyst displacement was small, we measured a mean displacement speed of 0.0125 +/− 0.0019 µm.min^-1^, which is in accordance with previous recording of wild-type cyst movements (Morris and Spradling, 2011).

**Figure 3:**
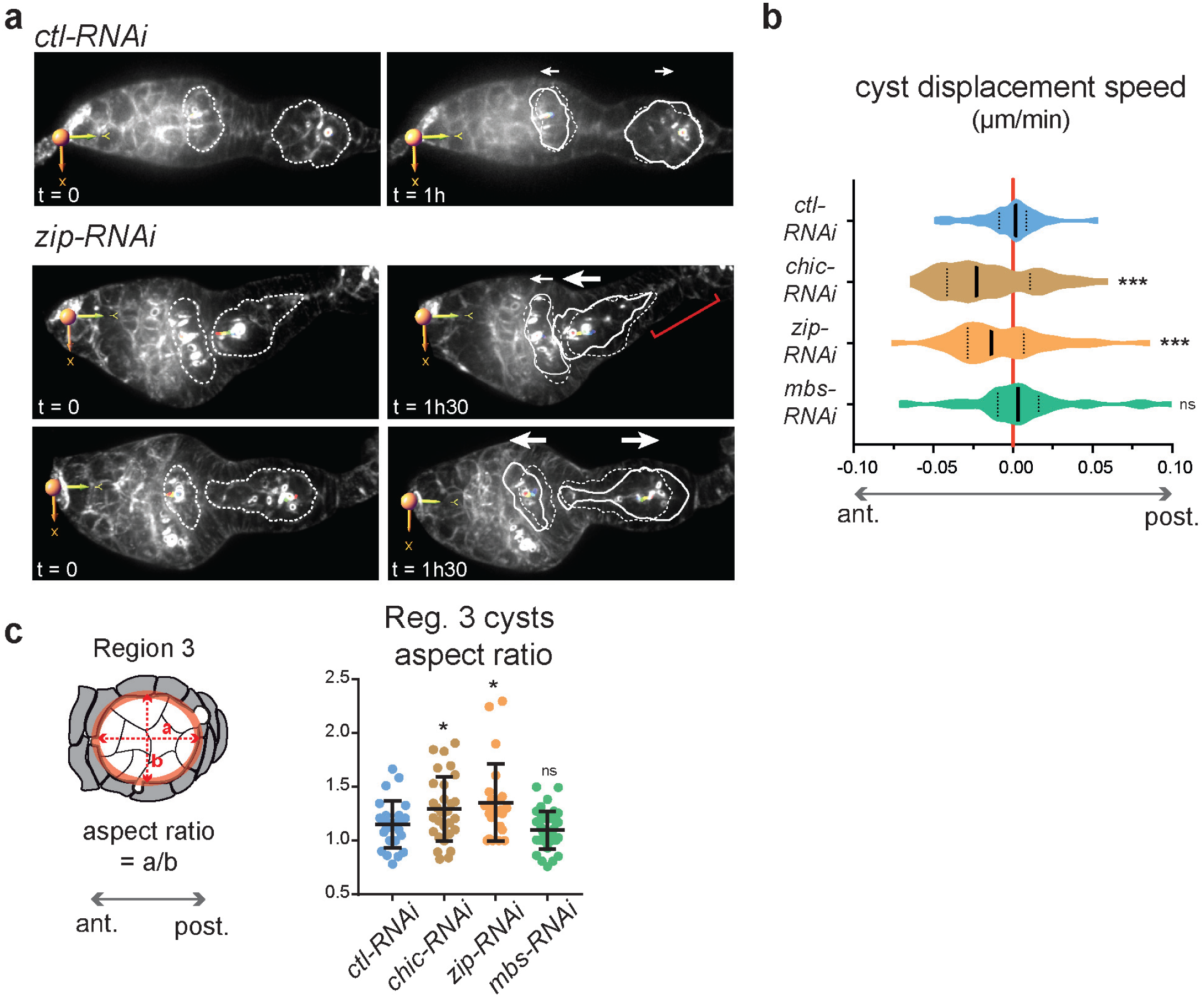
Alteration of germline contractility leads to defective sorting during encapsulation. (**a**) Still images from movies of *ctl-RNAi* and *zip-RNAi* germaria. Germline cysts were tracked over time. Cap cells at the anterior of the germarium serve as a reference point. Original cyst position is indicated with dashed lines and final cyst position is indicated with plain line. White arrows indicate cyst displacement. Red bracket indicates long stalk. (**b**) Quantification of cyst displacement speed along the a-p axis depending on the different mutant conditions. Violin plots with median and 25%-75% quartiles are shown. n = 44 *ctl-RNAi* cysts; n = 56 *chic-RNAi* cysts; n = 54 *zip-RNAi* cysts; n = 63 *mbs-RNAi* cysts. *** *P* < 0.001, ns, non-significant, Mann-Whitney *U-*test (performed on absolute speed values). (**c**, left) Schematic of a region 3 germline cyst surrounded by somatic cells. To measure the elongation of the cyst along the a-p axis, we measured the aspect ratio of the fitted ellipse (red) as illustrated. (right) Quantification of region 3 cysts aspect ratio depending on the different mutant conditions. Dot and whisker plots are shown. n = 26 *ctl-RNAi*, n = 30 *chic-RNAi,* n = 25 *zip-RNAi,* n = 38 *mbs-RNAi*. * *P* < 0.05, ns, non-significant, *t*-test. See also Supplementary Figure 3 and Movies 8-9

In *chic-RNAi* and *zip-RNAi* conditions, however, cysts movements were significantly increased and we measured a mean displacement speed of 0.0285 +/− 0.0022 µm.min^-1^ and 0.0237 +/− 0.0024 µm.min^-1^ respectively (Figure 3b, Figure S3). Cysts in regions 2b and 3 were frequently pushed back together more anteriorly (Figure 3a, *zip-RNAi* upper panel). These increased movements resulted in collisions between cysts in the germarium and formation of long stretches of FCs. If collisions were not resolved at the time of egg chamber individualization, this would lead to the formation of a compound egg chamber containing two cysts packaged together. Thus, cortical contractility is required to prevent uncontrolled germline cysts movement and collision between cyst at the time of encapsulation.

In *chic-RNAi* and *zip-RNAi*, we also noticed that cysts in region 3 instead of being round adopted an elongated shape along the a-p axis compared to controls (Figure 3c, increased aspect ratio). Live-imaging showed cysts being squeezed and sometimes cut into several parts by ingressing FCs (Figure 3a, *zip-RNAi* lower panel, Movie 8 and 9). Cysts splitting events would give rise to egg chambers with less than 16 germ cells. In addition, different parts of the split cysts could also be packaged with adjacent cysts generating egg chambers with more than 16 cells. These results indicated that cortical contractility also conferred stiffness to germline cysts, preventing them from being squeezed and cut by surrounding FCs.

In *mbs*-RNAi, cysts movements and speed were also increased compared to control conditions. We measured a mean displacement speed of 0.0203 +/− 0.0029 µm.min^-1^. Whereas in *zip*-RNAi and *chic*-RNAi cysts tend to be more frequently pushed toward the anterior, *mbs*-RNAi cysts tend to move most frequently toward the posterior (Figure 3b). This behavior was more pronounced for cysts in region 3, that significantly moved faster toward the posterior than *ctl-RNAi* in region 3 (Figure S3). In the strongest instances, we observed collisions in the posterior regions of the germarium between a fast moving cyst and an older cyst resulting in the encapsulation of the two cysts together (Movie 9). In contrast to *chic-RNAi* and *zip-RNAi*, we never observed cysts being split in *mbs-RNAi*. *mbs-RNAi* cysts were not squeezed but remained round in region 3 with an aspect ratio close to 1 (Figure 3c). These results suggest that increasing cortical contractility in *mbs-RNAi* favor faster displacement of the cysts toward the posterior of the germarium.

Together, our live-imaging experiments showed that cortical contractility is required for correct positioning of cysts at the time of encapsulation both to avoid collisions between cysts and formation of long stalks of FCs devoid of germ cells. Germline contractility is also required to maintain cyst integrity as a group of 16 cells and avoid cyst splitting. These observations helped explain our cell-lineage analysis. Thus, we concluded that alteration of cortical contractility in germ cells induced a loss of coordination between germline and somatic cells movements, leading to encapsulation of abnormal numbers of germ cells. Next, we looked for links between germ cells and somatic cells that could mediate this coordination.

### 5) Cadherin-based adhesion is required for correct encapsulation

During oogenesis, interactions between germ cells and somatic cells are mediated by E-Cadherin (E-Cad) homophilic interactions (Godt and Tepass, 1998; Gonzalez-Reyes and St Johnston, 1998). We also showed previously that adherens junctions are present between germ cells around ring canals (Figure 4a, Movie 10) (Fichelson et al., 2010). β-catenin (*armadillo*, *arm* in *Drosophila*) and α-catenin link E-Cadherin to the underlying actomyosin cortex (Peifer et al., 1993; Sarpal et al., 2012; White et al., 1998).

**Figure 4:**
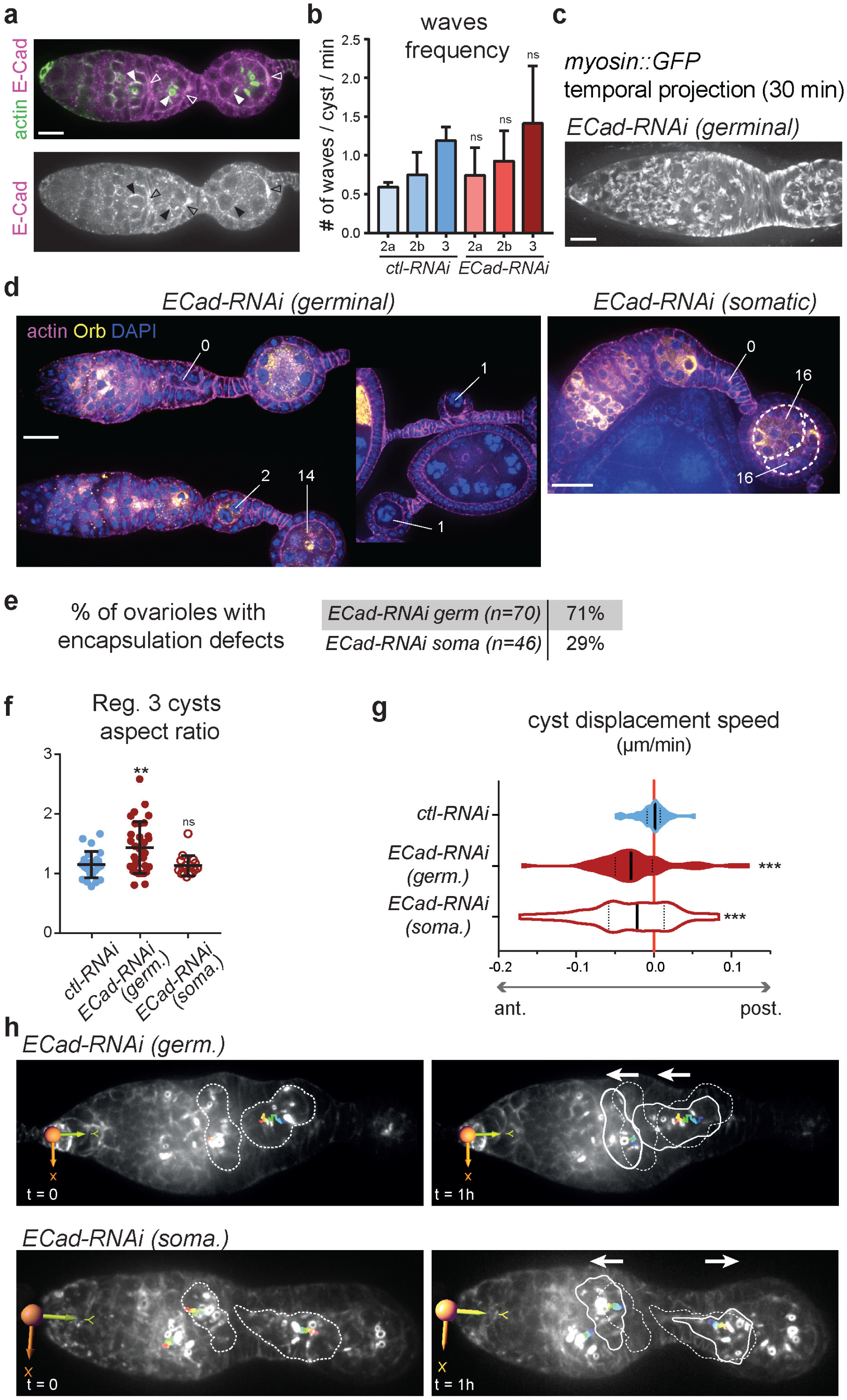
Cadherin-based adhesion is required for correct encapsulation. (**a**) Wild type germarium stained with E-Cad (magenta) and F-actin (green). E-Cad localized within germline cells around ring canals (plain arrows) and between germline cysts and the somatic layer (empty arrows). (**b**) Quantification of wave frequency per cyst in region 2 and 3 of the germarium. Mean and SD are shown. n = 6 cysts in region 2a, n = 8 cysts in region 2b, n = 6 cysts in region 3, 5 *ECad-RNAi* germaria. ns, not significant, *t*-test. (**c**) Temporal projection of a 30 min movie of a *ECad-RNAi* germarium expressing *sqh::GFP* to follow myosin waves. (**d**) Fixed images of ovarioles stained with phalloidin to mark actin (magenta), Orb that marks the oocyte (yellow) and DAPI that marks the DNA (blue). Depletion of *ECad* in the germline (*ECad-RNAi germinal*) or in the follicle cells (*Ecad-RNAi somatic*) lead to encapsulation defects and the formation of egg chambers with abnormal number of cerm cells. White dotted lines underline two different cysts packaged together in one egg chamber. Numbers indicate the number of cells included in each unit. (**e**) Quantification of the occurrence of encapsulation phenotypes in fixed ovariols when *Ecad* is depleted in the germline (*Ecad-RNAi germ.*) or in the somatic cells (*Ecad-RNAi soma.)*. (**f**) Quantification of region 3 cysts aspect ratio depending on the different mutant conditions. Dot and whisker plots are shown. n = 26 *ctl-RNAi*, n = 33 *ECad-RNAi germ,* n = 17 *ECad-RNAi soma.* ** *P* < 0.01, ns, not significant, *t*-test. (**g**) Quantification of cyst displacement speed along the a-p axis depending on the different mutant conditions. Violin plots with median and 25%-75% quartiles are shown. n = 53 *ECad-RNAi germ* cysts; n = 39 *ECad-RNAi soma* cysts. *** *P* < 0.001, Mann-Whitney *U-*test (performed on absolute speed values). (**h**) Still images from movies of *Ecad-RNAi germ* and *ECad-RNAi soma* germaria. Germline cysts were tracked over time. Cap cells at the anterior of the germarium serve as a non-moving reference point. Original cyst position is indicated with dashed lines whereas final cyst position is indicated with plain line. White arrows indicate cyst displacement Anterior is on the left and posterior on the right. Scale bars, 10μm (c), 20μm (a, d). See also Movies 10-13 and Supplementary Figure 4.

To decrease homophilic interactions between germ and somatic cells, we depleted E-Cad (*shotgun*, *shg* in Drosophila) either in the germline using the *bam*-Gal 4 driver (*ECad-RNAi germ*, Figure S3a) or in the FCs using *traffic jam*-Gal4, which is strongly expressed in FCs (*ECad-RNAi soma*, Figure S3a). We used E-Cad-shRNA to avoid functional compensation by N-Cad (see Material and Methods and (Loyer et al., 2015)). We found that it induced encapsulation phenotypes, similar to those observed after knocked-down of *zip* or *chic* in the germline. On fixed ovaries, we found egg chambers made of two 16-cell cysts, or separated by empty stalk cells (Figure 4d, e). In addition, when *E-Cad* was depleted in germ cells, we found a majority of egg chambers with fewer germ cells, indicating that groups of 16 cells had been split between several egg chambers (Figure 4d). As in *chic-RNAi* or *zip-RNAi*, occurrence of split cysts correlated with strong deformations of *ECad-RNAi germ* mutant cysts along the a-p axis in region 3 (Figure 4f). However, egg chambers containing split cyst were not observed when *E-Cad* was depleted in FCs only, and cyst aspect ratio in region 3 was not affected in this case (Figure 4f).

Live imaging confirmed these encapsulation defects: we observed increased cyst movements in region 2b and 3 of the germarium (mean displacement speed = 0.0427 +/− 0.0044 µm.min^-1^ for *ECad-RNAi germ* and 0.0507 +/− 0.0066 µm.min^-1^ for *ECad-RNAi soma*), with cysts being pushed toward the anterior or the posterior leading to collisions with the preceding or following cyst (Figure 4g, h, Figure S4b, Movie 11). We also observed cysts being deformed along the a-p axis and split by FCs when *E-Cad* was knock-downed in the germline (Movies 12). Similar phenotypes were observed although with a lower penetrance after knock-down of *arm* (*arm-RNAi*) in the germline (Figure S4c, Movie 12). Consistent with our results, encapsulation defects were also reported with mutant alleles of *β-catenin* and *α-catenin* (Peifer et al., 1993; Sarpal et al., 2012; White et al., 1998).

These results showed that reducing cell adhesion in either germ cells or somatic cells gave similar encapsulation phenotypes than impairing cortical contractility in germ cells. We thus tested whether reducing cell adhesion affected germline cells contractility. Quantification of actomyosin waves frequency showed that they were not affected in *ECad-RNAi* mutant cysts (Figure 4b, c, Movie 13). Eliminating cell adhesion thus does not noticeably impact germ cells contractility. Junctional complexes, however, are required to transmit contractile forces to neighboring cells (Martin et al., 2010).

We concluded that correct encapsulation requires generation of contractile forces by germline cysts and adhesion between germ cells to prevent cysts splitting. It also requires transmission of these contractile forces to follicle cells layer through *E-Cad* adhesion complexes to maintain cyst position and prevent collisions between cysts.

### 6) Altering blebs frequency leads to encapsulation defects

Since blebs are direct consequences of strong cortical contractions and in direct contact with FCs, we investigated whether blebs were also involved in encapsulation and cyst positioning. To manipulate blebs occurrences without directly affecting actomyosin contractility, we thought of modifying properties of the cortex by expressing two different mutant forms of Moesin in germ cells. Moesin is the sole ERM (Ezrin Radixin Moesin) protein in *Drosophila* and links the actomyosin cortex to the plasma membrane. Its activity is regulated by phosphorylation at T559 (Kunda et al., 2008). When we expressed a non-phosphorylatable form (*moe-TA::GFP*) in germ cells, known for its dominant-negative function, we noticed an increase in the average number of blebs per cyst (Figure 5a, b). This correlated with encapsulation defects and the formation of egg chambers with abnormal numbers of germ cells (Figure 5c, d). Cell lineage analysis further showed that GFP+ cysts expressing *moe-TA::GFP* could be found packaged with a wild-type cyst in a compound egg chamber (Figure 5c). We then looked by live-imaging how these defects emerged. Interestingly, we observed that *moe-TA::GFP* overexpressing cysts in region 3 moved more and faster (with a mean displacement speed of 0.0187 +/-0.0028) toward the posterior than control cysts (mean displacement speed = 0.0107 +/-0.0028) (Figure 5e). A highly blebbing cyst could sometimes contact and invade a posterior cyst leading to cysts collision (Figure 5f, Movies 14, 15).

**Figure 5:**
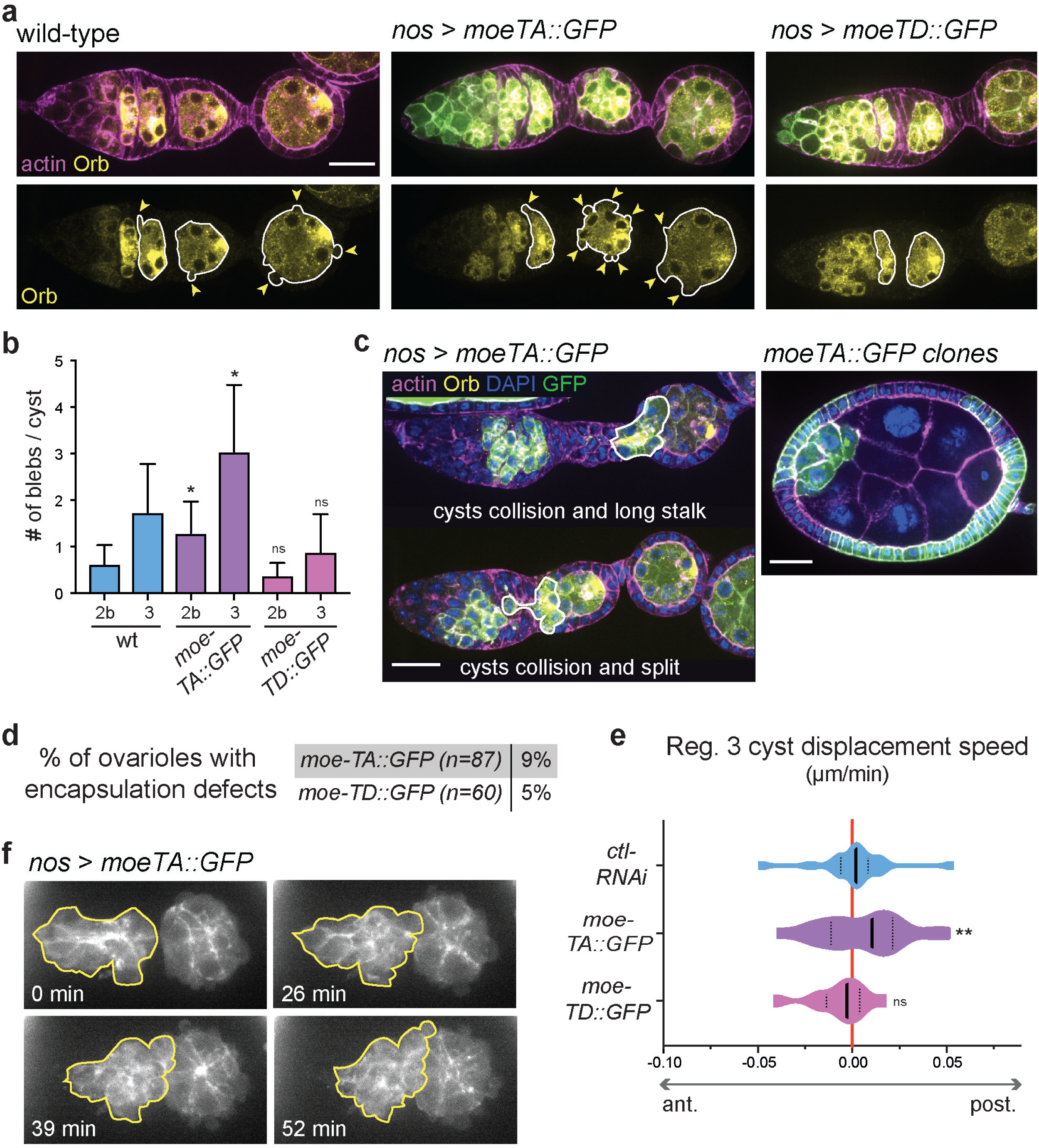
Increasing blebs frequency induces forward movements of germline cysts and cyst collisions. (**a**) wild-type germarium (left) and germaria overexpressing *moe-TA::GFP* or *moe-TD::GFP* in the germline (middle and left) stained with Orb (yellow) and phalloidin (actin, magenta). Cysts boundaries are underline in white. Bleb protrusions are indicated with arrows. (**b**) Quantification of the average number of blebs per cyst in region 2b and 3, depending on the different conditions. Mean and SD are shown. n = 7 cysts in region 2b, n = 9 cysts in region 3, 12 wild type germaria; n = 12 cysts in region 2b, n = 27 cysts in region 3, 20 *moe-TA::GFP* germaria, n = 8 cysts in region 2b, n = 11 cysts in region 3, 10 *moe-TD::GFP* germaria. * *P* < 0.05, ns, non-significant, *t*-test. (**c**) Fixed images of ovarioles stained with phalloidin to mark actin (magenta), Orb that marks the oocyte (yellow) and DAPI that marks the DNA (blue). (left) Overexpression of *moe-TA::GFP* in the germline lead to cyst collisions. The invading anterior cysts are underline in white. (Right) Compound egg chamber made of a *moe-TA::GFP* overexpressing cyst and one unmarked wild-type cyst. (**d**) Quantification of the occurrence of encapsulation phenotypes in fixed ovarioles with germinal overexpression of *moe-TA::GFP* or *moe-TD::GFP*. (**e**) Quantification of cyst displacement speed along the a-p axis depending on the different conditions. Only cysts in region 3 that have already transition from disc-shape to a rounder shape were considered. Violin plots with median and 25%-75% quartiles are shown. n = 21 cysts in region 3 for *moe-TA::GFP*; n = 12 cysts in region 3 for *moe-TD::GFP.* ns, non-significant. ** *P* < 0.01, Mann-Whitney *U-*test (performed on absolute speed values). (**f**) Still images from a movie of a germarium overexpressing *moe-TA::GFP* in the germline. An anterior cyst (yellow outline) migrate forward and collide with a posterior cyst. Scale bars, 20μm. See also Movies 14-15.

On the other hand, expressing a phosphomimetic form of Moesin (*moe-TD::GFP*) in germ cells is thought to induce a stiffer cortex. We observed smaller blebs and a decrease in number of blebs, although not statistically significant (Figure 5a, b). In contrast to cysts expressing *moe-TA::GFP*, cysts expressing *moe-TD::GFP* were not able to significantly move forward and only induced mild encapsulation defects (Figure 5d, e).

These results showed that blebs could play a role during encapsulation. Increasing bleb occurrences was sufficient to accelerate germline cysts movement, which can induce collisions and encapsulation defects. Thus, our results suggest that increasing blebs in germline cysts could induce a bleb-based motility behavior.

### 7) Germline cysts play an active role in cysts sorting using migration-like mechanisms

To reveal a putative germ cells autonomous role in cyst positioning during encapsulation, we aimed to block somatic cells movement and thus suppress constriction forces exerted on germline cysts. We did so by mechanically blocking FCs centripetal migration. We observed that when we mounted germarium in halocarbon oil (10S), FCs strongly adhere to the coverslip and were unable to migrate. In normal conditions, FCs convergent-extensions movements constrict the germarium in-between cysts progressively reducing the width of the stalk that will separate the future egg chambers (Figure 6a) (Morris and Spradling, 2011). In hydrogel we measured a reduction of 4.5 +/− 0.5 % of the stalk width in 50 min. In oil, FCs tend instead to slightly expand on the coverslip resulting in an expansion of 0.6 +/− 1.0 % of the stalk width (Figure 6a). FCs were thus not able to intercalate and constrict underlying germ cells. In these conditions, we measured positive movement of germline cysts toward the posterior of the germarium, indicating that wild-type germline cysts were able to migrate on stalled FCs, which can lead to collision between cysts (Figure 6b, c, Movie 16). Knock-down of *zip* or *E-Cad* in the germline however significantly reduced germline cyst movement and speed in germarium mounted in oil (Figure 6b). This indicated that the ability of cysts to migrate on stalled FCs depends on cortical contractility and adhesion with surrounding FCs. Importantly, these results also revealed that fast displacement of germline cysts with reduced cortical contractility or adhesion observed in hydrogel were passive and imposed by surrounding FCs constriction forces (Figure 3b and Figure 4g). Mutant cysts were passively pushed backward or forward as somatic cells rearranged and constricted to form a stalk (Figure S5).

**Figure 6:**
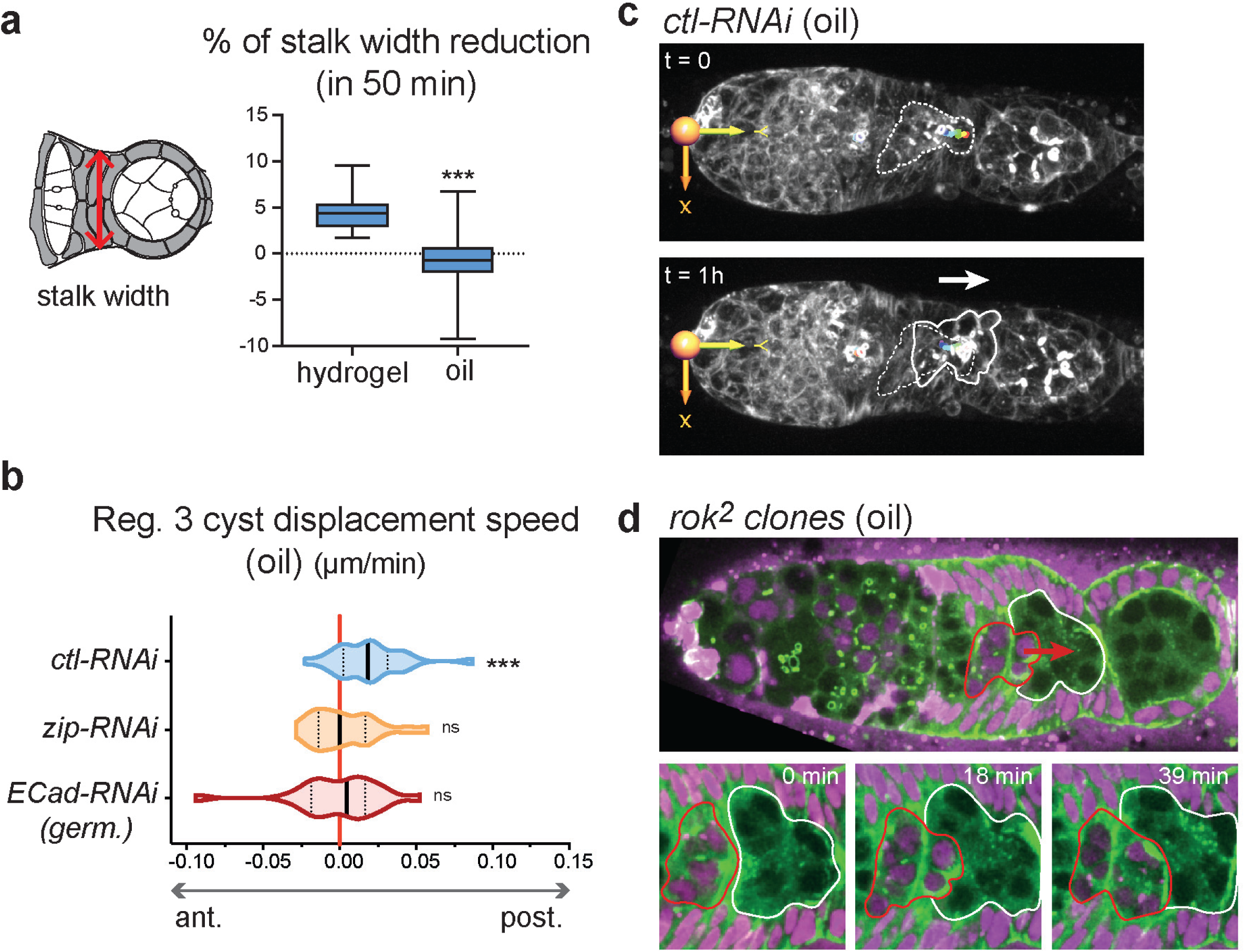
Germline cyst can migrate when somatic cell movement is blocked. (**a**, left) Schematic of the posterior part of the germarium. FCs form a stalk that separate the two cysts. Red arrow indicates the measure of the stalk width. (right) Quantification of the percentage of stalk width reduction during 50 min of recording, for germaria mounted either in hydrogel or in oil (negative values indicate expansion). Box and whisker plots are shown. n = 20 germaria mounted in hydrogel, n = 13 germaria mounted in oil. *** *P* < 0.001, *t*-test. (**b**) Quantification of cyst displacement speed along the a-p axis when germaria are mounted in oil. Violin plots with median and 25%-75% quartiles are shown. n=26 cyst in Region 3 ctl-RNAi; 25 cysts in region 3 for *zip-RNAi,* 22 cysts in region 3 for *shg-RNAi*. *** *P* < 0.001, ns, non-significant. One sample Wilcoxon signed rank test (test against the null hypothesis m = 0, no displacement). (**c**) Still images from a movie of a control germarium (*ctl-RNAi*) mounted in oil. An anterior cyst in region 3 migrate forward and collides with a posterior cyst. Original cyst position is indicated with dashed lines whereas final cyst position is indicated with plain line. White arrows indicate cyst displacement. (**d**) Still images of a mosaic germarium containing wild-type (RFP+, magenta) germline cyst and *rok^2^* mutant cysts. (bottom) High magnification are shown. A wild-type cyst (RFP+, red outline) migrates forward and invade the position of a mutant cyst (unmarked, white outline). Red arrow indicates cyst displacement. See also Movies 16-17 and Supplementary Figure 5.

Thus, our data suggest that cortical contractility in germ cells can generate forces enabling cysts to migrate. To support this idea, we compared two populations of germline cysts within the same germarium using clonal analysis. We induced mosaic germaria containing germline cysts mutant for *rok* and marked by the absence of RFP (wild-type cysts are RFP+) (Figure 6d). *rok* mutant cysts showed a clear reduction in waves frequency compared to wild type cysts (Figure S2a). In these conditions, wild type cysts invaded into posterior *rok* mutant cysts, indicating that wild type cysts were consistently faster to the posterior than *rok* mutant cysts (Figure 6d, Movie 17). Consistently, we never detected the reverse invasion of wild type cysts by mutant cysts (n=6 invasions of *rok^2^* mutant cysts by wild-type cysts, 9 mosaic germaria).

Altogether, these results suggest that in normal conditions, germline cysts use migration-like mechanisms to maintain their position within the germarium during encapsulation. Their autonomous movement toward the posterior is usually masked by the prominent movements and constriction of FCs around them that tend to push neighboring cysts in opposite direction to separate them. However, if we mechanically block FCs movements, germline cysts can visibly migrate forward (Figure 7).

**Figure 7:**
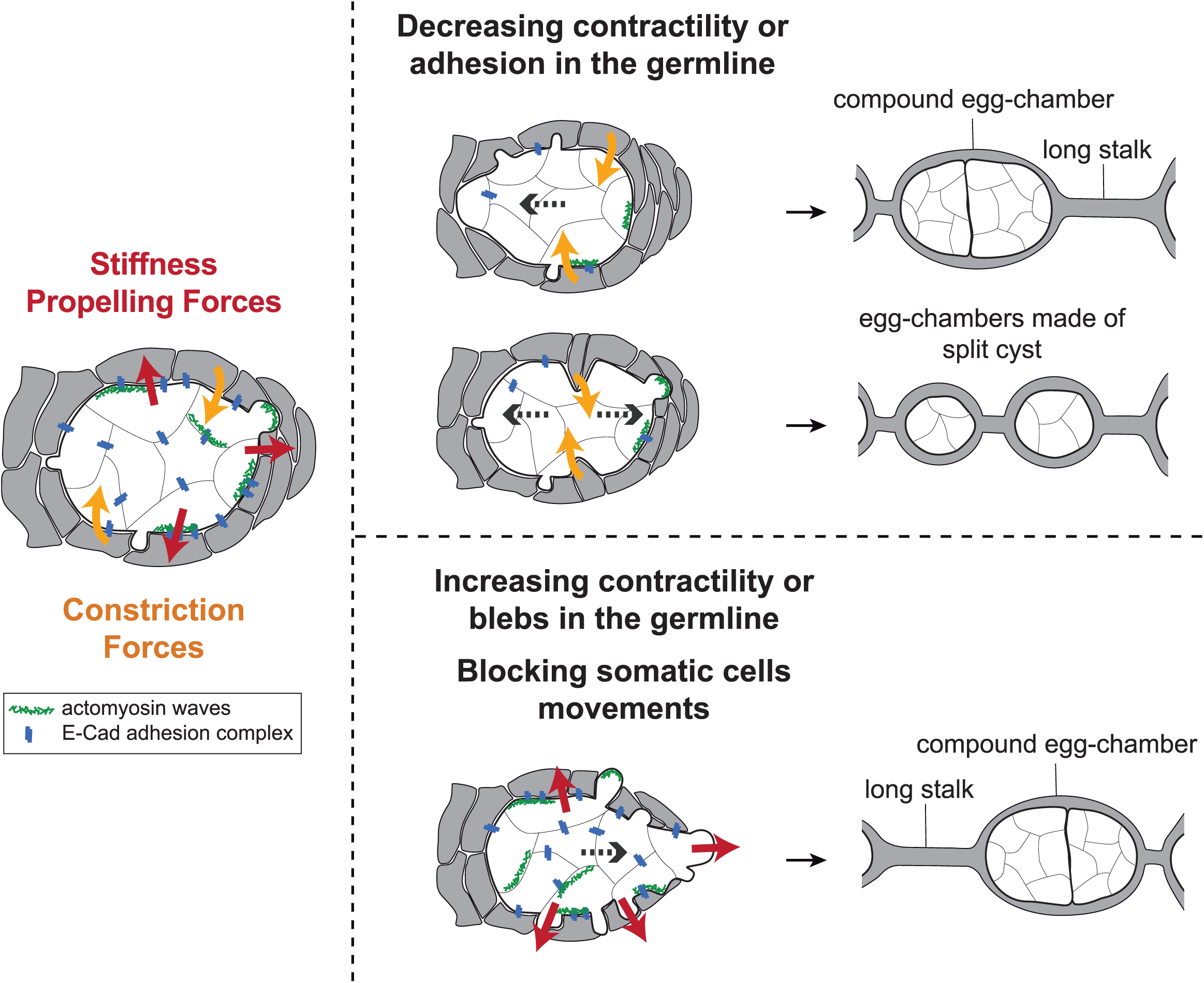
Encapsulation requires a proper balance between germline and somatic forces. (left) Germline cysts exert mechanical forces dependent on cortical contractility and adhesion (red arrows). This confers stiffness within the 16-cell cyst to maintain cyst integrity and propelling forces to maintain proper cyst positioning during encapsulation. At the same time, somatic cells migrate around germline cyst exerting constriction forces (orange arrows). In wild type, germline forces resist constriction forces exerted by surrounding somatic cells and maintain correct cyst position during encapsulation. (middle) Modifying the equilibrium between germline and somatic forces decouples cysts movement from somatic cells movement. On one hand, decreasing germline contractility or adhesion induce germline cysts sliding backward or forward, leading to cysts collision, which can lead to the formation of compound egg chambers. decreasing germline contractility or adhesion also induce cyst splitting by constricting somatic cells, which generate incomplete egg chambers. On another hand, increasing germline contractility or blebbing or blocking somatic cells movement can induces a net forward migration of the germline cysts that can also result in cyst collision and encapsulation defects.

## DISCUSSION

Our study revealed the existence of periodic contractile waves of the actomyosin network at the cortex of germ cells, as described in other developmental systems (Bement et al., 2015; Maître et al., 2015; Weiner et al., 2007). The nature of these waves is dual, requiring both motor activity and actin polymerization. We demonstrated that these contractions are required to maintain correct positioning of germline cysts during encapsulation by FCs and to maintain their integrity as groups of 16 cells. In light of our results, we propose that there are at least two kinds of forces at play during encapsulation (Figure 7). Convergent-intercalation of somatic cells exerts constriction forces on the underlying germline cells, while contractility and protrusions of germ cells exert propelling forces on the overlying FCs. Correct encapsulation requires a proper balance between these forces. Indeed, on the one hand, when germ cells contractility was weakened, somatic cells could squeeze and cut or displace germline cysts. On the other hand, when somatic cells convergence was blocked, germline cells migrated faster and collided. We further showed that E-Cadherin-based adhesion between germline and somatic cells is required to transmit forces and to coordinate both morphogenetic movements.

Mechanistically, we propose that contractile waves contribute to the dynamics of cellular adhesion between germline and somatic cells. Indeed, knocking down contractility in germ cells or cadherin-based adhesion between germline cysts and FCs both led to increased and uncontrolled movements of germline cysts. This resulted in encapsulation defects with the packaging of several cysts in the same egg chamber, or long stalk of FCs without germ cells. We propose that cortical contractility at the interface between germline cyst and FCs helps to remodel dynamically cellular adhesion and thus to coordinate germline and somatic cells morphogenetic movements. This could involve junctional strengthening under tension or contractility-dependent E-Cad turnover and junction remodeling (Cavey et al., 2008; Liu et al., 2010; Shigenobu et al., 2010). In addition, friction forces could be generated between germline and somatic cells by flows of actomyosin linked to transmembrane E-cad proteins. This mechanism would be reminiscent of single cell migration mechanisms used in confined or crowded environments (Bergert et al., 2015; Hawkins et al., 2011; Paluch et al., 2016).

Reducing contractility or adhesion specifically in germ cells induced an additional phenotype whereby germline cysts were deformed along the anterior-posterior axis and split by ingressing somatic cells. This phenotype was not seen (or very rarely) when removing cadherin in somatic cells only. It indicates that adhesion and contractility in germ cells are required to sort cysts into group of 16 cells and to resist constriction forces applied by somatic cells. Adhesion and contractility are commonly assumed to function in cell sorting by controlling tissue surface tension (Krens and Heisenberg, 2011; Maitre et al., 2012; Manning et al., 2010). We propose that a similar mechanism operates here, where contractility and adhesion between cells within a cyst favor the formation of a stiff sphere that cannot be split during encapsulation. Interestingly, the abnormal packaging of germ cells in flies with reduced adhesion or contractility in the germline, is similar to the normal aggregation of germ cells from different cysts into nests that occurs in mice oogenesis (Lei and Spradling, 2013). Our results suggest that simple differences in the regulation of cellular adhesion or cortical contractility could explain this evolutionary difference between mice and flies.

We also showed that germline cysts are blebbing during encapsulation. The importance of blebs for cell migration has been clearly demonstrated in several cases (Liu et al., 2015; Paluch and Raz, 2013; Paluch et al., 2006; Ruprecht et al., 2015). The underlying mechanisms and physical models, however, vary. One model postulates that blebs could engage in cell-cell adhesion with its environment and allow the forward transfer of cytoplasm. It has also been proposed that in a confined environment, blebs could push laterally allowing cells “to chimney” their way forward. Our results are compatible with a role of blebs in generating propelling forces during encapsulation. We found that increasing blebs frequency can induce posterior cyst migration. The underlying physical model remains, however, to be investigated. Together, our results suggest that germline cysts use migration-like mechanism such as blebbing and DE-Cad based friction with the somatic layer to maintain their position in the germarium during encapsulation, counterbalancing the forces exerted by ingressing FCs.

The reverse situation of somatic follicle cells migrating in between germ cells has been well characterized in later stages of *Drosophila* oogenesis. A small group of anterior follicle cells, called border cells (BCs), detach from the anterior pole of the egg chamber and migrate between nurse cells to reach the oocyte (Montell, 2003). Like encapsulation, this process is collective and requires DE-Cadherin, actin polymerization and myosin activity. However, despite these similarities, BCs follow a mesenchymal-like mode of migration by extending long protrusions toward the oocyte (Mishra et al., 2019). In contrast, we found that germ cells contractility induces the formation of blebs, which are associated with amoeboid migration (Lammermann and Sixt, 2009). Our results are more reminiscent of primordial germ cells migration described in zebrafish, indicating that blebs formation may be a conserved property of germline cells (Blaser et al., 2006).

Our results are a first step toward a mechanical understanding of encapsulation. Mechanical signals can instruct and pattern cell behaviors (Chanet and Martin, 2014). In light of our study, we can speculate that mechanical feedbacks between germ cells and somatic cells could play a role in self-organizing encapsulation. An important issue is to address how these mechanical inputs are integrated and regulated by biochemical signaling pathways. In both mice and flies, there is a wealth of literature describing cell-cell communications between somatic cells, and between germline and somatic cells at these stages. In *Drosophila*, disruptions of the Hedgehog, Wingless, Notch, Jak/Stat or EGF pathways, all lead to encapsulation defects with the formation of compound egg chambers with multiple cysts or long stalks devoid of germ cells (Bastock and St Johnston, 2008; Klusza and Deng, 2011; Roth and Lynch, 2009). These phenotypes have been attributed mostly to defects in cell fate specifications. In light of our study, these signaling pathways could also have a more direct role and regulate actomyosin contractility or adhesion, as their disruption induce similar defects. It will be exciting to re-analyze these biochemical and mechanical activities as well as their interplay during encapsulation. An integrated model of encapsulation will clearly benefit our understanding of gamete formation and reproductive biology.

## Acknowledgments

We are grateful to Juliette Mathieu for the initial observations of germline cysts being split in *shg* RNAi. We thank A. Guichet, A.C. Martin, A. Royou, the Bloomington Drosophila Stock Center and the Developmental Studies Hybridoma Bank for kindly providing flies and antibodies used in this study. We thank all members of the J-R. H. lab, F. Schweisguth, M-E. Terret and M. Malartre for discussions and helpful comments on the manuscript. S.C. is supported by an ARC postdoc fellowship and work in JRH lab is supported by CNRS, Inserm, Collège de France, FRM (Equipe FRM DEQ20160334884), ANR (ANR-15-CE13-0001-01, AbsCyStem) and Bettencourt-Schueller foundations.

## MATERIALS and METHODS

### Fly stocks and genetics

Myosin was visualized in live germaria using myosin regulatory light chain (*sqh* in *Drosophila*) fused to GFP, sqh::GFP (Royou et al., 2002) or mCherry, sqh::mCherry (Martin et al., 2008) expressed under its own promotor. Actin was visualized using the F-actin binding domain of Utrophin fused to GFP (Utr::GFP) expressed under an ubiquitous promotor (*sqh* promotor, (Rauzi et al., 2010)) or using the UASp-LifeAct::GFP (Huelsmann et al., 2013) construct express in the germline using the nanos–GAL4-VP16 (nos–GAL4, (Van Doren et al., 1998) driver. To visualize ROCK in live, we used a wild-type ROCK allele fused to GFP expressed under an ubiquitous promotor, ubip–GFP::ROCK (Bardet et al., 2013). To visualize DE-Cad in live, we used a knock-in insertion of GFP at the DE-Cad locus (Huang et al., 2009), and to visualize Armadillo we used the stock arm::GFP (BDSC#8556). Microtubules were visualized in live using the microtubule-associated protein Jupiter fused to GFP, Jup::GFP (Morin et al., 2001). Photoconversion experiments were performed using the stock w; sqh::Dendra2 (Roubinet et al., 2017).

For knockdown experiments, the following stocks were used: v;; UASp–white-shRNA (BDSC #35573), v;; UASp–chic-shRNA (BDSC #34523), v; UASp–zip-shRNA (BDSC #37480), v; UASp–mbs-shRNA (BDSC #41625), v; UASp–shg-shRNA (BDSC #38207), v; UASp–arm-shRNA (BDSC #35004). For this study, we generated: sqh::GFP; UASp–chic-shRNA, w; UASp–zip-shRNA; Utr::GFP, w; UASp–mbs-shRNA; Utr::GFP using stocks previously described.

Germline clones mutant for *shg^R69^* or *shg^IG29^*, which are null alleles of *shg*, rarely induced encapsulation defects in contrast to *shg*-shRNA. We believed it is caused by the functional compensation between E-Cad and N-Cad in the *Drosophila* germline as published previously for multinucleation phenotypes (Loyer et al., 2015). As shown by Loyet et al., *shg*-shRNA does not trigger compensation by N-Cad, we thus used this combination to disrupt cellular adhesion. In addition, we obtained identical encapsulation phenotypes with *arm*-shRNA, which links both E-Cad and N-Cad to the actomyosin cytoskeleton.

The white-shRNA was used as a control (*ctl-RNAi*) because white is not expressed during oogenesis. Depending on the strength of the shRNA, different drivers were used for knockdowns in the germline. To knockdown *zip* and *mbs* we used nos-GAL4, either the original stock or w; sqh::GFP; nos-GAL4 (generated using stocks previously described). To knockdown *chic* and *shg* we used a bam–GAL4(x2) driver (containing two copies of the bam-GAL4 driver, (Clemot et al., 2018)): w; bam–GAL4(x2) or w; bam–GAL4(x2); Utr::GFP (generated using stocks previously described). For gene knockdowns in follicle cells, we used: w; Traffic jam–GAL4; Utr::GFP (generated using stocks previously described and from Bloomington). Knock downs were performed at 29°C to increase the efficiency of the GAL4 driver.

To generate Flp out clones we used the stock: w hs-Flp; UASp–GFP; act–FRT-y+-FRT-GAL4 (generated using stocks from Bloomington). Heat-shocks were performed on early pupae, 30 min at 37°C.

*rok^2^* and *shq^1^* clones were generated using the Flp/FRT technique. The following stocks were used: w FRT-19A rok^2^ (Winter et al., 2001), w hs-Flp FRT-19A ubi–mRFP.nls and w FRT^101^ sqh^1^ (Karess et al., 1991), w hs-Flp FRT^101^ ubi–GFP. To induce clones, heat-shocks were performed on L2 larvae for 1h at 37°C for two consecutive days.

To modify cortex properties, favoring or reducing blebs occurrences, we expressed tagged version of Moesin in the germline using nos-GAL4 driver. w; UASp–moe-TA::GFP, w; UASp–moe-TD::GFP are gifts from A. Guichet. We also used: w; UASp–moe-TA::GFP; Utr::GFP and w; UASp–moe-TA::GFP; Utr::GFP (generated using stocks previously described). Crosses were done at 29°C to increase the efficiency of the GAL4 driver.

### Live and fixed imaging

5-day-old females were collected and dissected for live imaging or fixed experiments. *Live imaging in hydrogel* was adapted from (Burnett et al., 2018). Ovaries were dissected in Schneider medium (Sigma-Aldrich), and transfer onto a round 25 mm coverslip. The coverslips were previously coated with 3-(trimethoxysilyl)propyl methacrylate (Sigma-Aldrich). Medium was removed and 15 μL of 10% PEG-DA hydrogel solution (esibio) with 0.1% I2959 (photo initiator, Sigma-Aldrich) was added on the samples. A coverslip treated with deperlent was placed over the hydrogel droplet and the coverslip/coverslip sandwich was then placed over a UV light source and illuminated for 30s at 312nm for gelation. The upper coverslip was removed and the coverslip supporting the hydrogel disc was then placed into a chamber (Chamlide) filled with Schneider medium. See Supplemental Figure 1b. All imaging was performed at 25°C.

*Drugs treatment*. Few microliters of chemical or vehicle were added directly to the culture chamber while imaging. Cyto-D (Enzo Life Sciences) and CK-666 (Sigma) were diluted in DMSO (Sigma). Final concentrations in Schneider medium after addition of the drugs to the culture chamber were 2.5μM for Cyto-D and 50μM for CK-666. Y27632 (Enzo Life Sciences) was diluted in water and added to the chamber for a final concentration of 500μM. Colcemid (Sigma) was added for a final concentration of 62μg.mL^-1^.

*For live imaging in oil*, ovaries were dissected in oil (10S, Voltalef, VWR) and transfer onto a coverslip. Germaria were made to stick to the coverslip in oil.

*For immunostaining*, ovaries were dissected in PBS, fixed in 4%PFA, permeabilized in PBT (0.2%Triton) for 30 min, left overnight with primary antibodies in PBT at 4°C, washed 3 times 30 min in PBT, left with secondary antibody for 2 h at room temperature, washed 3 times 30 min in PBT and mounted in Cityfluor. The following primary antibodies were used: Orb (Mouse, 1:500, Developmental Studies Hybridoma Bank, DSHB, 4H8) and DE-Cadherin2 (Rat, 1:50, DSMB, Hybridoma Product DCAD2). Secondary antibodies used were Cy3 and Cy5 (1:200, Jackson laboratories). AlexaFluor568 phalloidin (Invitrogen) was used to visualize F-actin (1:400), and DAPI (Invitrogen) was used at 1:200.

All images were acquired on an inverted spinning-disc confocal microscope (Roper/Nikon) operated by Metamorph 7.7 coupled to a sCMOS camera and with a 60X/1.4 oil objective.

### Image processing and analysis

Images were processed using Fiji and Imaris (Bitplane) and graphs were generated in Prism (GraphPad). A bleach correction was applied to time-laps images. Images of the movies represent a maximum intensity Z projection (15μm). Waves and blebs frequency (Figure 1, 2 and 3) were measured in Fiji, every occurrence of a wave or bleb was counted, the resulting number was then divided by the time of the recording. Wave occurrences were either counted per cell (Figure 1f) or per cysts (Figures 1i, 2b, 3b). To measure the average number of blebs per cyst (Figure 5), we counted the number of blebs per cyst for each time frame (every 30 sec) and divided by the number of frames. This method was used to take into account bleb persistence.

Region 3 cyst aspect ratios (Figure 3) were measured in Fiji using the build-in toolbox, an ellipse was fitted to the shape of the cyst (as indicated in Figure 3f).

Cyst displacement were tracked in Imaris. Cap cells at the anterior tip of the germarium were used as a fixed reference point. To estimate cyst displacement, we tracked the oldest ring canal of each cyst, which is the widest and brightest, making it an easy object to track. We projected displacement along the a-p axis using Imaris build-in toolbox and calculated the mean speed over 1 to 2h. By convention, we conferred a negative value for displacement speed toward the anterior and a positive value for displacement speed toward the posterior.

To calculate the percentage of stalk width reduction over 50 min in hydrogel *vs.* in oil, we measure the width of the stalk at t = 0 and at t = 50 min and divided the difference by the initial width.

### Quantification and statistical analyses

Statistical analyses were performed using the Prism (Graphpad) statistics toolbox. For waves frequency and aspect ratio, *P* values were calculated using an unpaired *t*-test, the reference sample is the distribution in wild-type. To compare cysts displacements speed in our different conditions, we used a non-parametric Mann-Whitney *U-*test. the reference sample is the distribution in *ctl-RNAi*. Statistical analyses were performed on absolute value of cyst displacement speed. To compare constriction of the stalk in hydrogel vs. oil, *P* value was calculated using an unpaired *t*-test, the reference sample is the distribution in hydrogel. Cysts autonomous migration on stalled FCs (Figure 6b) was tested using a non-parametric one sample Wilcoxon test. *P* values were calculated against the null hypothesis m = 0 (no displacement).

---

**Supplementary Figure 1. Related to Figure 1.**
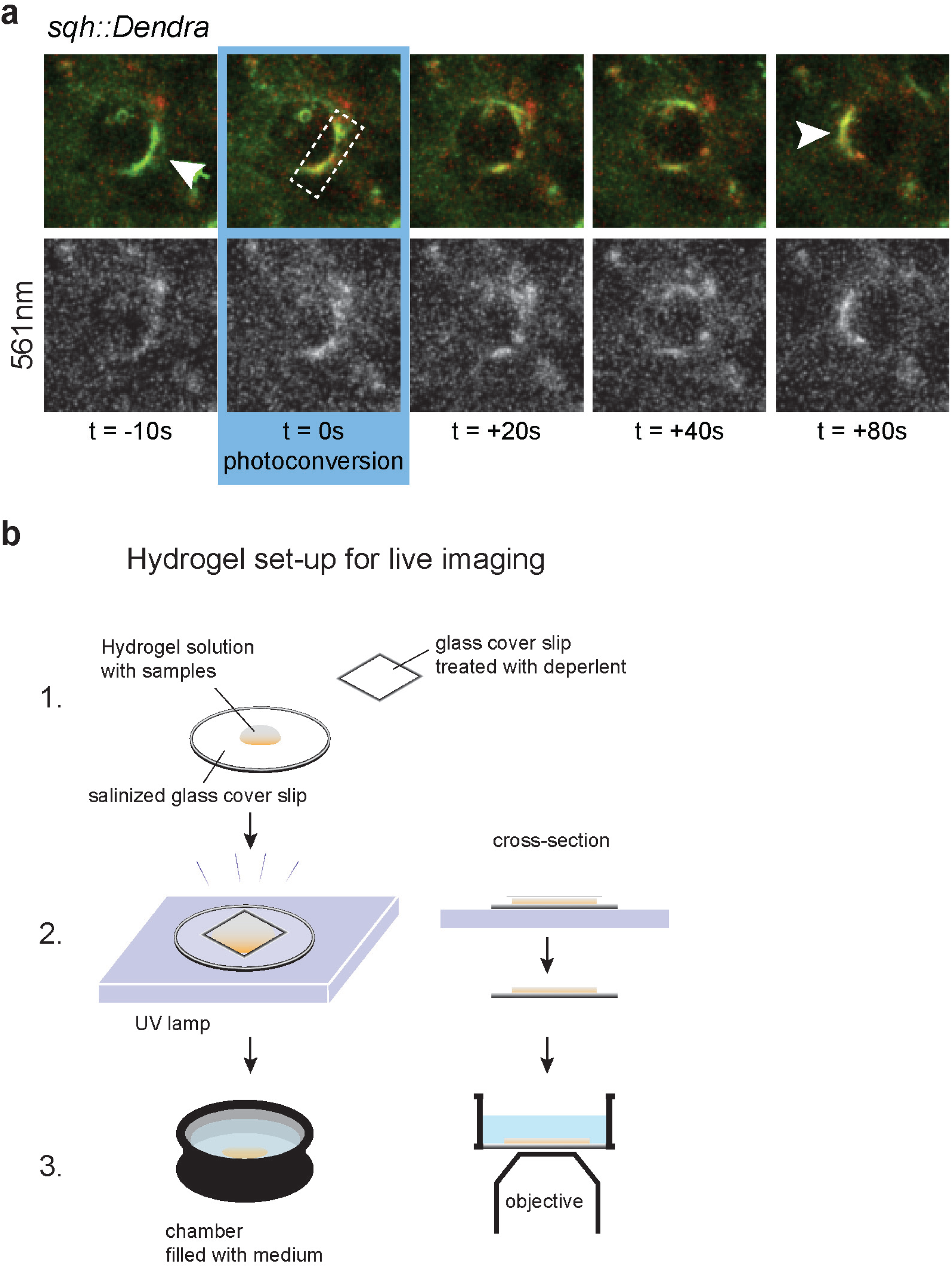
(**a**) photoconversion experiments reveled traveling waves. Time-laps images of a germ cell expressing *sqh::Dendra2* that stains myosin. *sqh::Dendra2* molecules were photoconverted with blue light (405 nm) at t = 0 within the indicated ROI (dashed line). Bottom panels show converted molecules (revealed with 561nm laser). (**b**) Schematic of the hydrogel set-up for live imaging. See Methods.

**Supplementary Figure 2. Related to Figure 2.**
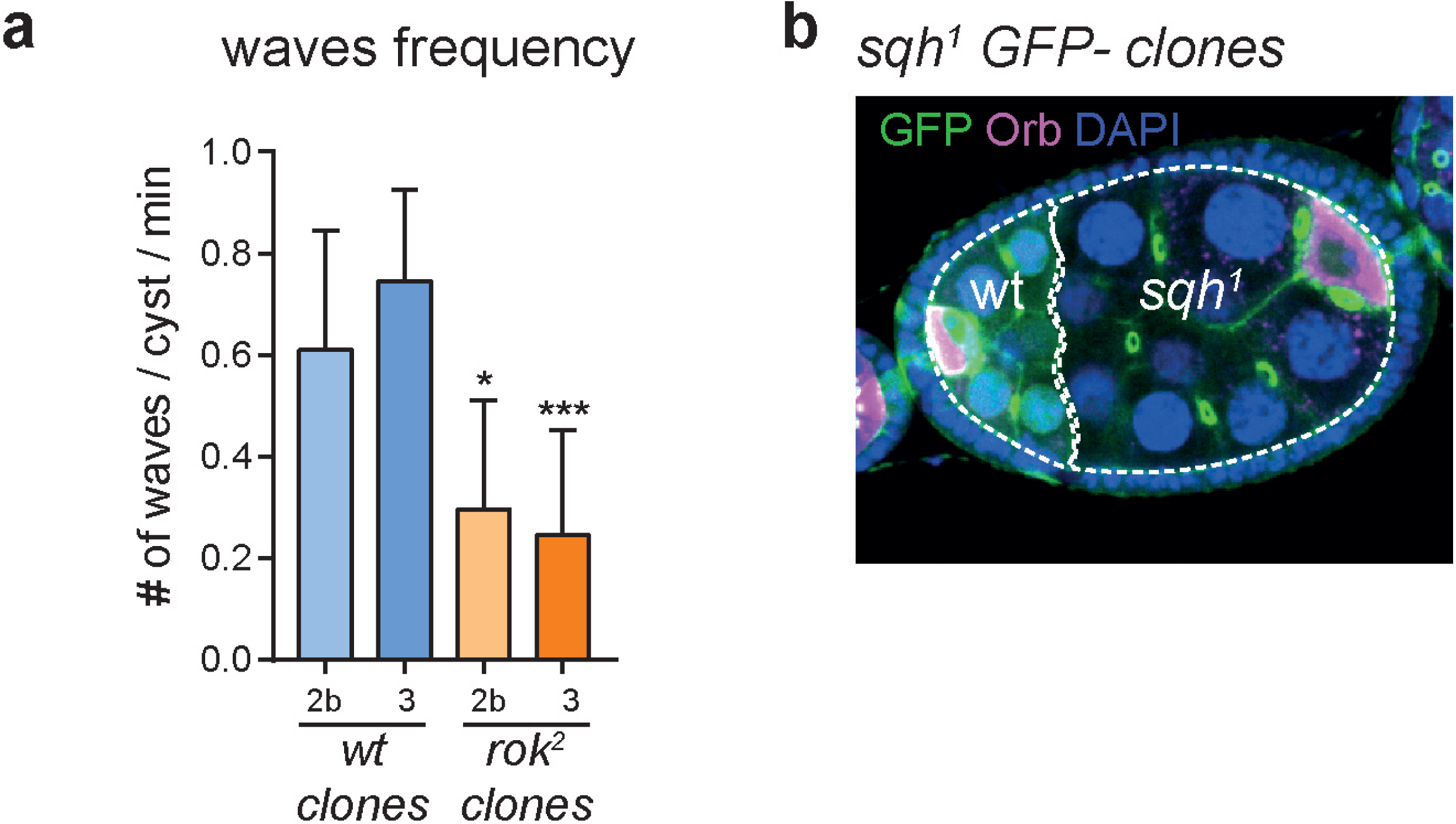
(**a**) Quantification of wave frequency per cyst in region 2b and 3 of the germarium of wild type and *rok^2^* clones. Mean and SD are shown. n = 10 wt cysts in region 2b, n = 6 wt cysts in region 3; n = 6 *rok^2^* cysts in region 2b, n = 5 *rok^2^* cysts in region 3; 14 mosaic germaria. * *P* < 0.05, *** *P* < 0.001, *t*-test. (**b**) Fixed image stained with Orb that marks oocytes (magenta) and DAPI that marks the DNA (blue) of a compound egg chamber containing a wild type (GFP+) and a *sqh^1^* mutant cyst. White dotted line underlines the two cysts packaged together.

**Supplementary Figure 3. Related to Figure 3.**
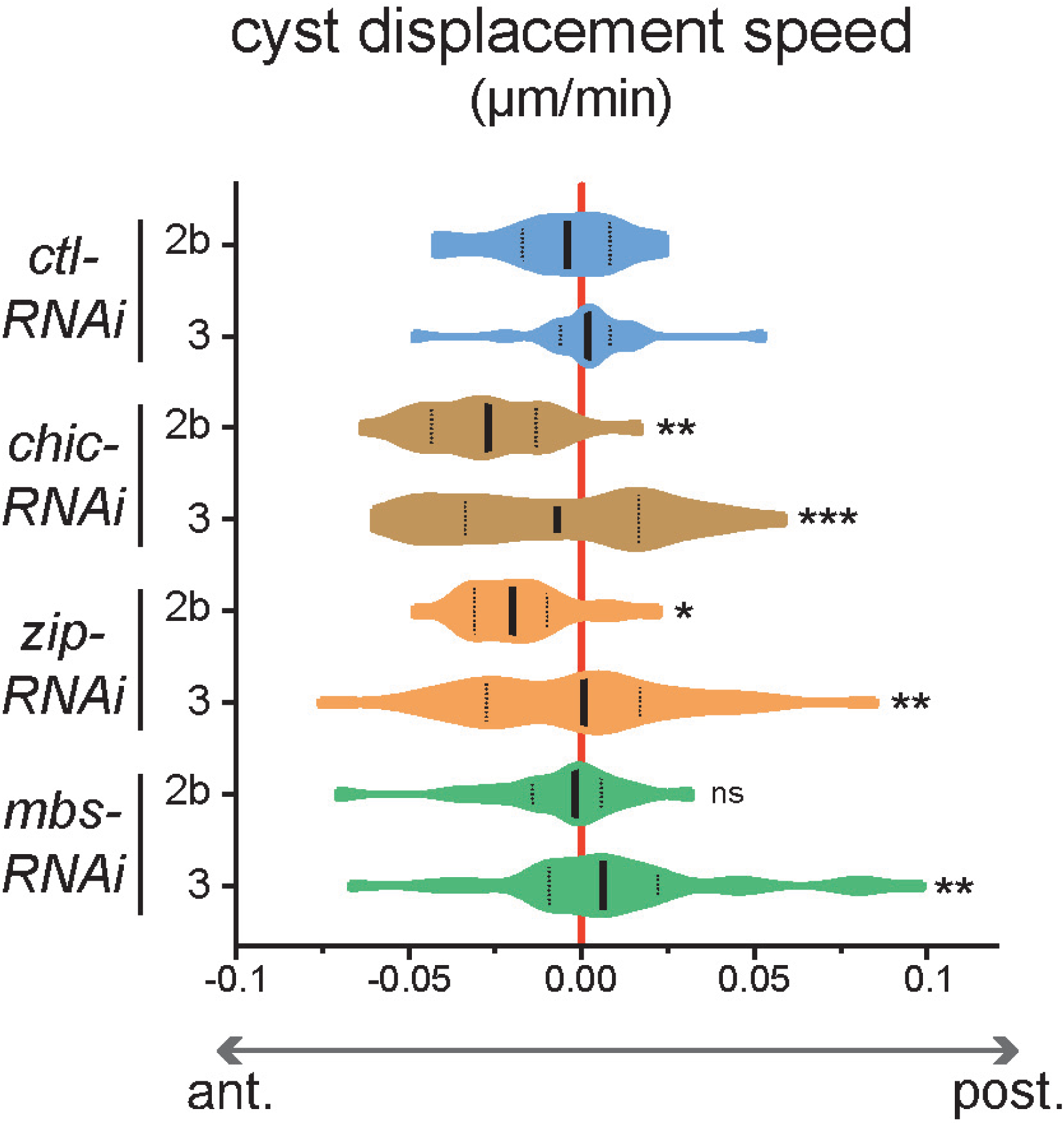
Quantification of cyst displacement speed along the a-p axis for the different indicated mutant conditions. The figure represents the same data set as in Figure 3b with the detailed displacement speed for cysts either in region 2b (positioned roughly in the middle of the germarium and with a disc shape) or in region 3 (positioned at the posterior of the germarium and with a rounder shape). Violin plots with median and 25%-75% quartiles are shown. n = 20 cysts in region 2b, 24 cysts in region 3 for *ctl-RNAi*; n = 25 cysts in region 2b, 31 cysts in region 3 for *chic-RNAi*; n = 21 cysts in region 2b, 33 cysts in region 3 for *zip-RNAi*; n = 21 cysts in region 2b, 42 cysts in region 3 for *mbs-RNAi*. * *P* < 0.05, ** *P* < 0.01, *** *P* < 0.001, ns, non-significant, Mann-Whitney *U-*test (performed on absolute speed values).

**Supplementary Figure 4. Related to Figure 4.**
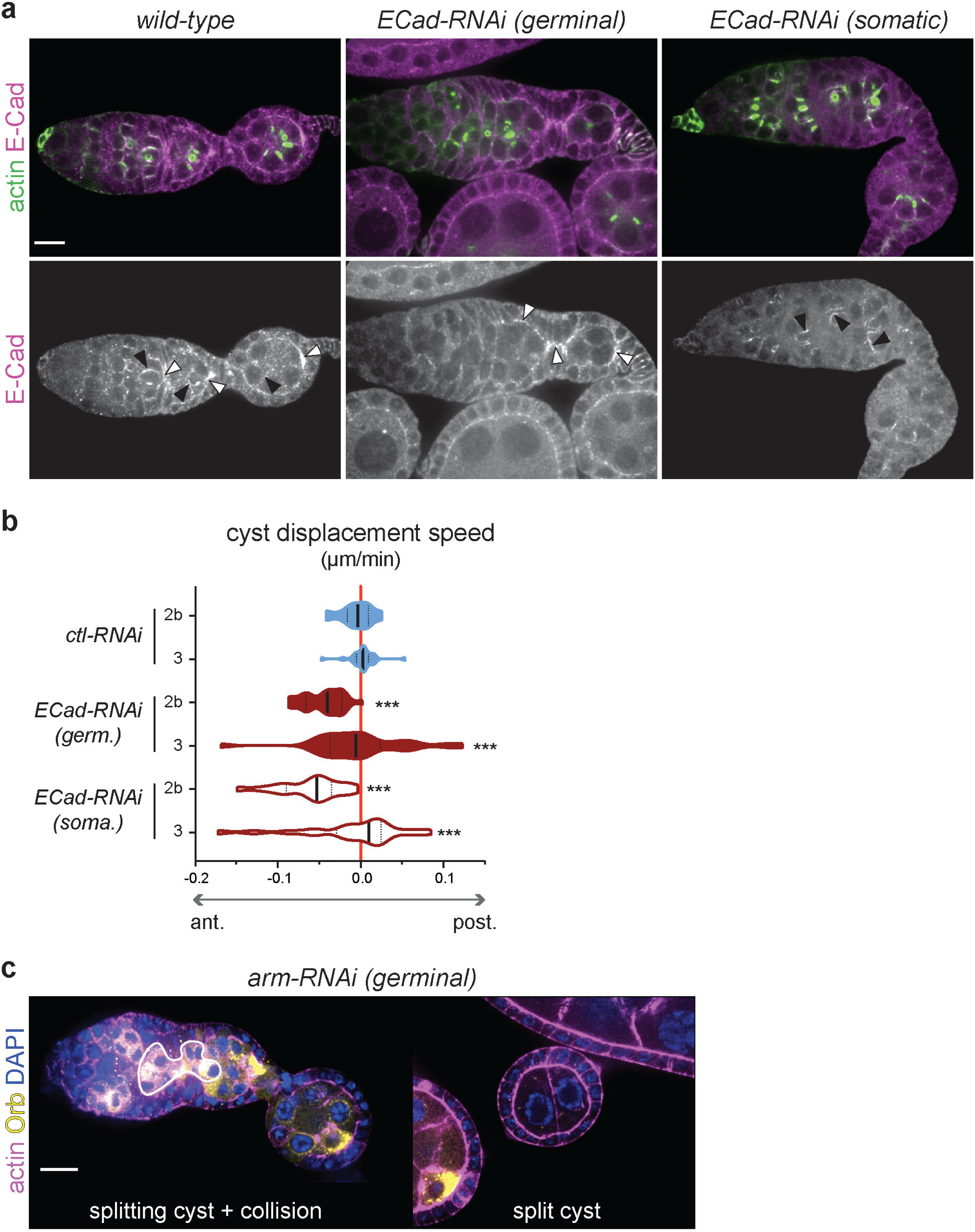
(**a**) Fixed germaria expressing *Utr::GFP* (actin, green) and stained for E-Cad. (right) E-Cad localized within germline cells around ring canals (plain arrows) and between germline cysts and the somatic layer in wild type (empty arrows). (middle) germline knock down of *ECad* specifically reduce E-Cad staining between germ cells. (left) somatic knock down of *ECad* specifically reduce E-Cad staining between somatic cells. (**b**) Quantification of cyst displacement speed along the a-p axis depending on the different mutant conditions. The figure represents the same data set as in Figure 4f with the detailed displacement speed for cysts either in region 2b (positioned roughly in the middle of the germarium and with a disc shape) or in region 3 (positioned at the posterior of the germarium and with a rounder shape). Violin plots with median and 25%-75% quartiles are shown. n = 25 cysts in region 2b, 28 cysts in region 3 for *shg-RNAi germ.*; n = 17 cysts in region 2b, 22 cysts in region 3 for *shg-RNAi soma.* *** *P* < 0.001, Mann-Whitney *U-*test (performed on absolute speed values). (**c**) Fixed images of ovarioles stained with phalloidin to mark actin (magenta), Orb that marks the oocyte (yellow) and DAPI that marks the DNA (blue). Germline knock down of *arm* leads to encapsulation defects similar to germline knock down of *shg*, such as cyst collision within the germarium and cyst splitting. Scale bars, 20μm

**Supplementary Figure 5. Related to Figure 6.**
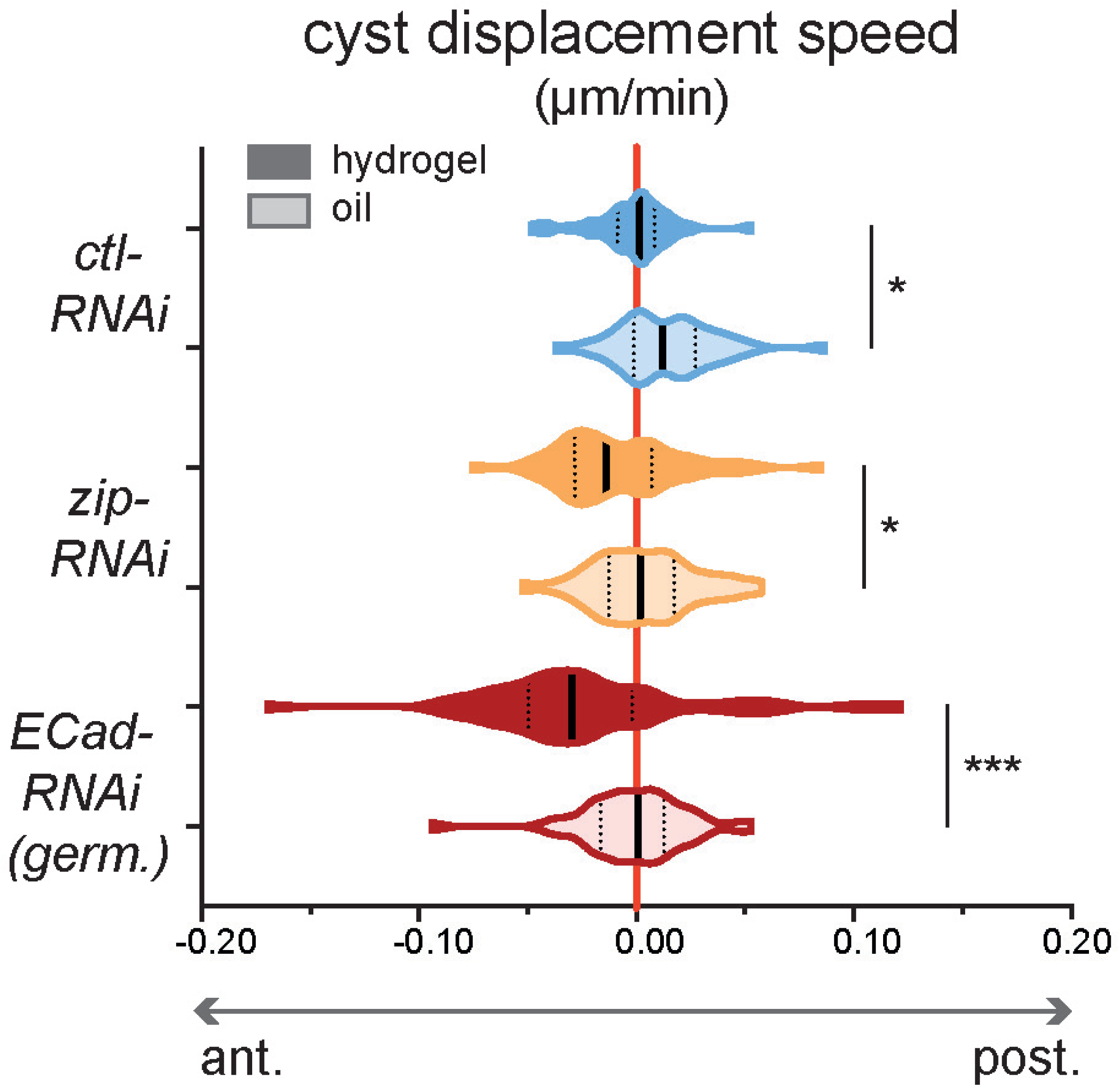
Comparison of cyst displacement speed along the a-p axis when samples are mounted in hydrogel or in oil for the different conditions as indicated. Violin plots with median and 25%-75% quartiles are shown. For measurements in hydrogel, n = 44 *ctl-RNAi* cysts; n = 54 *zip-RNAi* cysts; n = 53 *ECad-RNAi* cysts. For measurements in oil, n = 44 *ctl-RNAi* cysts; n = 43 *zip-RNAi* cysts; n = 46 *ECad-RNAi* cysts. * *P* < 0.05, *** *P* < 0.001, Mann-Whitnay *U-*test.

## MOVIES LEGENDS

**Movie 1: Cortical myosin waves in germ cells.** Related to Figure 1.

Time-lapse images from a germarium expressing the myosin marker sqh::GFP.

**Movie 2: Cortical F-actin waves in germ cells.** Related to Figure 1.

Time-lapse images from a germarium expressing the F-actin marker LifeAct::GFP in the germline (using the nos-GAL4 driver). The contour of the germarium is outlined.

**Movie 3: Blocking myosin activity or actin polymerization using chemical inhibitors stops wave’s propagation.** Related to Figure 1.

Time-lapse images from germaria expressing the myosin marker sqh::GFP or the F-actin marker Utr::GFP as indicated. Drugs or vehicle (DMSO) were added to the culture medium at 6 min as indicated.

**Movie 4: Depolymerizing microtubule does not affect wave’s propagation.** Related to Figure 1.

Time-lapse images from germaria expressing the microtubule marker Jup::GFP (top) or the myosin marker sqh::GFP (bottom) as indicated. Colcemid was added to the culture medium at 6 min as indicated. Addition of Colcemid immediately depolymerize microtubules but does not affect myosin waves.

**Movie 5: Germline cysts are blebbing.** Related to Figure 1.

Time-lapse images from a germarium expressing the myosin marker sqh::GFP and cytoplasmic GFP in the germline (using the nos-GAL4 driver). Arrows indicate blebs protrusion.

**Movie 6: Inhibiting cortical contractility by adding Cyto-D immediately stops blebs**. Related to Figure 1.

Time-lapse images from a germarium expressing the myosin marker sqh::GFP and cytoplasmic GFP in the germline (using the nos-GAL4 driver). Cyto-D was added to the culture medium at 15min.

**Movie 7: Genetic modifications of cortical contraction waves.** Related to Figure 2.

Time-lapse images from germaria expressing the myosin marker sqh::GFP and the indicated RNAi in the germline. *chic-RNAi* and *zip-RNAi* reduce the number of waves per cyst, whereas *mbs-RNAi* increases the number of waves per cyst.

**Movie 8: Reducing germline contractility affects cyst displacement during encapsulation.** Related to Figure 3.

Time-lapse images from germaria expressing the F-actin marker Utr::GFP and the indicated RNAi in the germline. Cysts displacements along the a-p (y) axis were tracked over time. A reference point was position at the anterior end of the germaria were cap cells reside. The displacement track is time-colored from blue to red.

**Movie 9: Modifying germline contractility can lead to cyst splitting or collision.** Related to Figure 3.

Time-lapse images from germaria expressing the F-actin marker Utr::GFP and the indicated RNAi in the germline. (top) Example of a split cyst in *chic-RNAi*. (bottom) Example of a collision in *mbs-RNAi*.

**Movie 10: Dynamic junctional complexes between germ cells and between germ cells and somatic cells.** Related to Figure 4.

Time-lapse images from germaria expressing the junctional markers ECad::GFP (top) or arm::GFP (bottom).

**Movie 11: Reducing adhesion affects cyst displacement during encapsulation.** Related to Figure 4.

Time-lapse images from germaria expressing the F-actin marker Utr::GFP and *shg-RNAi* either in the germline (top) or in the somatic cells (bottom). Cysts displacements along the a-p (y) axis were tracked over time. A reference point was position at the anterior end of the germaria were cap cells reside. The displacement track is time-colored from blue to red.

**Movie 12: Decreasing germ cells adhesion can lead to cyst splitting.** Related to Figure 4.

Time-lapse images from germaria expressing the F-actin marker Utr::GFP and the indicated RNAi in the germline. Two examples of cysts being split (arrows) are shown, in *shg-RNAi* (top) and in *arm-RNAi* (bottom).

**Movie 13: Decreasing DE-Cad in the germline does not affect cortical waves.** Related to Figure 4.

Time-lapse images from germaria expressing the myosin marker sqh::GFP and the indicated RNAi in the germline.

**Movie 14: Increasing blebs frequency can induce forward cyst movement and cysts collision.** Related to Figure 5.

(top) Time-lapse images from a germarium expressing moe-TA::GFP in the germline (using the nos-GAL4 driver) and the F-actin marker Utr::GFP. Cyst displacement along the a-p (y) axis were tracked over time. A reference point was position at the anterior end of the germarium were cap cells reside. The displacement track is time-colored from blue to red. (bottom) Example of a collision. Time-lapse images from a germarium expressing moe-TA::GFP in the germline (using the nos-GAL4 driver). An anterior cyst migrates forward and invade the position of a more posterior cyst.

**Movie 15: Increasing blebs frequency can induce cysts collision.** Related to Figure 5.

Time-lapse images from a germarium expressing moe-TA::GFP in the germline (using the nos-GAL4 driver). A higly blebbing cyst invade the position of a more posterior cyst. This germarium was mounted in oil.

**Movie 16: Mechanically blocking somatic cells movements can induce forward cysts migration.** Related to Figure 6.

Time-lapse images from a germarium expressing the F-actin marker Utr::GFP and the a *ctl-RNAi* in the germline. Cysts displacements along the a-p (y) axis were tracked over time. A reference point was position at the anterior end of the germarium were cap cells reside. The displacement track is time-colored from blue to red. The germarium was mounted in oil that blocks somatic cell centripetal migration.

**Movie 17: wild-type cysts migrate faster than rok2 mutant cysts.** Related to Figure 6.

Time-lapse images from a mosaic germarium containing wild-type (RFP+) germline cysts and *rok^2^* mutant cysts (marked by the absence of RFP), and expressing the myosin marker sqh::GFP. A wild-type cyst (asterisk) migrate and invade the position of a more posterior mutant cyst. This germarium was mounted in oil.

